# Scalable high-fidelity human vascularized cortical assembloids recapitulate neurovascular co-development and cell specialization

**DOI:** 10.64898/2026.01.09.698753

**Authors:** Shubhang Bhalla, Belda Gulsuyu, Damian Sanchez, Jayden M. Ross, Santhosh Arul, Adnan Gopinadhan, Muhammet Öztürk, Tanzila Mukhtar, Jonathan J. Augustin, Jerry C. Wang, Joseph Kim, Chang N. Kim, Sena Oten, Yohei Rosen, John M. Bernabei, Vijay Letchuman, Shantel Weinsheimer, Helen Kim, Elizabeth E. Crouch, Edward F. Chang, David Haussler, Mircea Teodorescu, Arnold R. Kriegstein, Tomasz J. Nowakowski, Ethan A. Winkler

## Abstract

Human cortical development involves the coupling of neurogenesis and cerebrovascular growth. However, interactions between neural and vascular cells are largely missing in most brain organoids, which are crucial models for studying neurodevelopment and disease. Here, we establish vascularized cortical assembloids (vCAs) by fusing mesoderm-derived vascular organoids (VOs) with cortical organoids (COs). vCAs self-assemble lumenized networks of endothelial cells, pericytes, and perivascular fibroblasts that acquire blood–brain barrier (BBB) specialization and arteriovenous specification in vitro. Single-cell RNA-sequencing and immunofluorescence imaging revealed that vascularization improves neuroepithelial architecture, reduces hypoxia and apoptosis, expands cortical progenitor pools, and enhances neuronal maturation and connectivity compared with COs. Moreover, atlas-level integration with human neurodevelopmental tissue and cross-protocol benchmarking demonstrate superior transcriptional concordance, especially for vascular cell and glial populations, relative to existing approaches. This scalable and reproducible platform improves fidelity and throughput for modeling human neurovascular co-development and enables systematic studies across brain regions and diseases using engineered or patient-derived iPSCs.

## Introduction

Human cortical development proceeds through coordinated neural progenitor proliferation, migration, and differentiation, which is influenced by concurrent cerebrovascular growth and specialization^1-5^. Cortical vascularization and neurogenesis progress in tandem to build neural circuits and expand the cortex^4,6^, with bidirectional neurovascular, trophic, and paracrine cues regulating the production and patterning of new neurons and vascular specialization^7-10^. Developing blood vessels are composed of endothelial cells, pericytes, smooth muscle cells, and perivascular fibroblasts^11,12^, and early establishment of this neurovascular unit couples glia and neurons to the vasculature to shape angioarchitecture and confer specialized properties, such as the blood-brain barrier (BBB)^9,13-15^. Yet, the mechanisms governing human neurovascular co-development remain difficult to study at scale, and current approaches often rely on two-dimensional cell culture, animal models, or post-mortem tissues, each lacking human-specific tissue complexity or cellular dynamics.

Organoids offer a potential avenue to study cortical vascularization and neurodevelopment, but the distinct developmental origins of neural and vascular lineages have hindered their development *^6,16^*. Cortical organoids recapitulate cardinal features of human corticogenesis, including ventricular/subventricular progenitor zones, temporal differentiation patterns, and early lamination^17-20^, and they are vital for studying otherwise inaccessible neurodevelopmental milestones. However, widely used differentiation protocols exclude mesoderm-derived cell lineages, including vascular and immune populations^17,18^. Without vasculature, organoids lose vascular trophic cues and rely on diffusion for nourishment, which limits growth and induces hypoxia and metabolic stress. These differences may restrict neuronal maturation and subtype specification, ultimately reducing fidelity to the human brain^21,22^. Emerging organoid vascularization strategies remain limited in fidelity or scale^16^. For example, rodent xenotransplantation of cortical organoids introduces host vasculature but sacrifices human specificity and throughput^23,24^. Endothelial co-cultures form vessel-like structures with limited perivascular cell diversity^25-27^. Vascular-neural fusion assembloids improve cell diversity, but often show asymmetric integration, non-physiological vessel geometry, limited throughput or reproducibility, and minimal benchmarking to primary human tissues has been performed ^28,29^.

Here, we develop a modular assembloid strategy that vascularizes cortical organoids with high-fidelity at scale. We generate mesoderm-derived vascular organoids containing endothelial cells, pericytes, and perivascular fibroblasts in 96-well format, and fuse them to cortical organoids in prefabricated molds to achieve symmetric vascular integration and reproducibility. This platform reconstitutes core neurovascular features of human cortical development, including BBB specialization, endothelial arteriovenous specification, and a supportive neural progenitor niche, while advancing neuronal maturation and connectivity entirely *in vitro*. Atlas-level integration with primary human tissues and cross-protocol benchmarking show that this model most faithfully recapitulates human neurovascular cortical co-development. Importantly, this modular vascularization method may be integrated with region-specific organoids and engineered using patient-derived induced pluripotent stem cells, enabling systematic studies of human cortical development and associated diseases.

## Results

### Vascularized cortical assembloids recapitulate key cellular features of human cortical development

During embryonic neurodevelopment, brain vasculogenesis begins with the formation of a mesoderm-derived perineural vascular plexus^4,6^. To recapitulate this process, we generated mesoderm-derived vascular organoids (VOs) and adapted vascular differentiation to a high-throughput 96-well format (Methods, **Fig. 1a**). Specifically, VOs were generated from human induced pluripotent stem cells (hiPSCs) through sequential mesodermal induction (days 0-3), vascular differentiation (days 3-6), and vascular sprouting in a 3D collagen I-Matrigel matrix (days 6-11) (**Fig. 1a**)^20,30,31^. By day 16, we detected platelet endothelial cell adhesion molecule (CD31^+^) endothelial cells organized into vascular networks, platelet-derived growth factor receptor-beta (PDGFRb^+^) pericytes, lumican (LUM^+^) perivascular fibroblasts, and allograft inflammatory factor 1 (IBA1^+^) perivascular myeloid cells (**Fig. 1b**). CD31^+^ endothelial sprouts extended radially into the collagen I-matrigel matrix, forming a self-organizing vascular plexus (**Extended Data Fig. 1a**). In parallel, we separately generated cortical organoids (COs) from hiPSCs with established neural induction protocols ^18,32^. COs progressed with expected temporal sequence of neural and glial differentiation, including SRY-Box Transcription Factor 2 (SOX2^+^) neural progenitor cells (NPC), homeodomain-only protein (HOPX^+^) outer radial glia cells, and eomesodermin homolog (EOMES^+^) intermediate progenitor cells by day 35, COUP-TF interacting protein 2 (CTIP2^+^) deep-layer and SATB homeobox 2 (SATB2^+^) upper layer cortical neurons by day 63, and glial fibrillary acidic protein (GFAP^+^), S100 calcium binding protein B (S100B^+^), aquaporin-4 (AQP4^+^) astrocytes by day 70 (**Fig. 1c**).

**Fig. 1.**
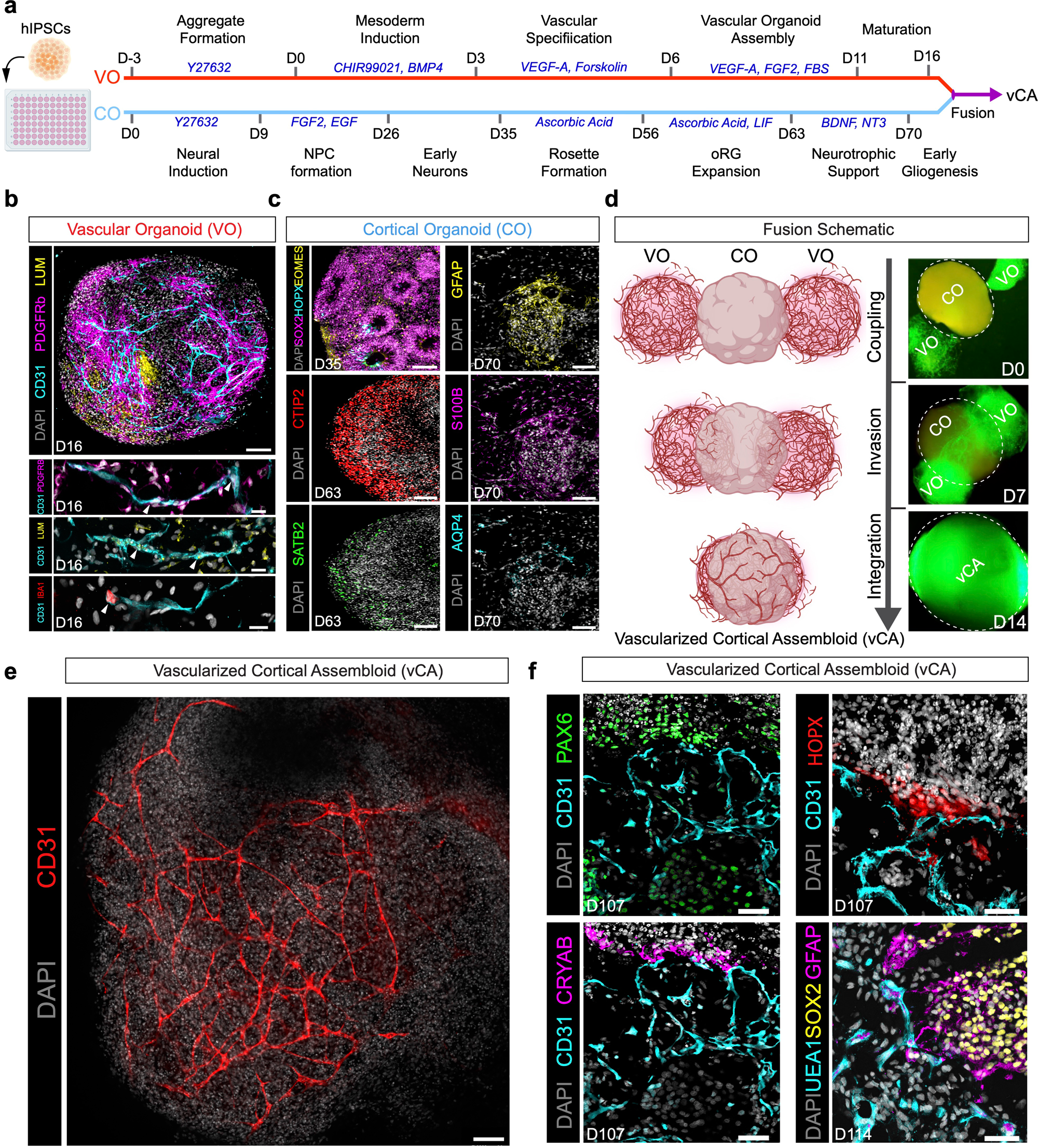
Modular generation and fusion of vascularized cortical assembloids. **a**, Schematic of parallel differentiation protocols for vascular organoids (VOs) and cortical organoids (COs) from hiPSCs. Red arrow, timeline for creation of VOs, including mesodermal induction (D3 to D0), vascular specification (D3 to D6), and vascular organoid assembly (D6 to D11). Blue arrow, timeline for creation of COs from neural induction (D0 to D9) to cortical maturation (D35 to D70) for COs. Key transcription factors and morphogenetic events indicated at respective timepoints. **b**, Representative confocal microscopy analysis of day 16 VOs showing CD31^+^ endothelial networks (cyan), PDGFRβ^+^ pericytes (magenta), LUM^+^ perivascular fibroblasts (yellow), and IBA1^+^ myeloid cells. Scale bars 100 μm for lower magnification image (top), 10 µm for high magnification images (bottom). **c**, Representative confocal microscopy analysis of COs showing temporal progression of cortical differentiation in COs. At day 35, SOX2^+^ (magenta), HOPX^+^ (cyan), and EOMES^+^ (yellow) progenitors were observed. Scale bar, 100 µm. At day 63, CTIP2^+^ deep-layer (red) and SATB2^+^ upper-layer neurons (green) were observed. Scale bars, 100 µm. At day 70, GFAP^+^ (yellow), S100B^+^ (magenta), and AQP4^+^ astrocytes were observed. Scale bars, 100 μm. **d**, Left, schematic showing creation of vascular cortical assembloids (vCAs). Right, Representative time-lapse, whole-mount microscopy analysis of GFP^+^ vascular organoid showing initial coupling (day 0), vascular invasion (day 7), and integration (day 14) following fusion with CO. **e**, Representative confocal microscopy analysis of day 100 vCAs showing CD31^+^ endothelial vascular networks. DAPI (white) labels cell nuclei. Scale bar, 100 μm. **f**, Representative confocal microscopy analysis of day 107 vCAs showing CD31^+^ endothelial cells (cyan), PAX6^+^ neural progenitors(green), CRYAB^+^ truncated radial glia (magenta), and HOPX^+^ outer radial glia (red). Bottom right, representative confocal microscopy analysis of day 114 vCAs showing UEA1^+^ endothelial cells (cyan), GFAP^+^ astrocytes (magneta), and SOX2^+^ neural progenitors (yellow). Scale bars, 50 μm.

During human neurodevelopment, the perineural vascular plexus forms next to the neural tube’s pial surface, and vascular cells invade neural tissue via sprouting angiogenesis^4,6^. To model this, we generated vascularized cortical assembloids (vCAs) by fusing two sprouting day-22 VOs to opposite poles of a day-72 CO in a single Matrigel droplet using prefabricated molds to maintain throughput and reproducibility while promoting symmetric vascularization (Methods, **Fig. 1d**). Using GFP-expressing hiPSCs, we tracked vascularization in real-time. GFP^+^ VO sprouts invaded COs by day 7 and consolidated into integrated GFP^+^ vascular networks by day 14 (**Fig. 1d**). In day-14 post-fusion assembloids, CD31 immunostaining showed robust, interconnected endothelial vascular networks throughout (**Fig. 1e**). Across vCAs from three hiPSC cell lines, vascular branching index, density, and length did not statistically vary, supporting the method’s reproducibility (**Extended Data Fig. 1b-c**). Compared to primary human cortical plate tissue samples (gestational week 22; n = 3 donors), vCAs showed no significant differences in CD31^+^ vascular branching index, density, or length (**Extended Data Fig. 1b-c**), confirming that vCAs closely recapitulate the angioarchitecture of the developing human cortex.

Because vascular-progenitor interactions shape many neurodevelopmental processes^7,9,10,33-36^, we next assessed whether vCAs re-establish known spatial relationships between the vasculature and neural/glial progenitors. CD31^+^ vessels localized adjacent to PAX6^+^ NPCs (**Fig. 1f**, top left) and HOPX^+^ outer radial glia (**Fig.1f**, top right), supporting close physical proximity between the vasculature and relevant progenitor populations. CD31^+^ endothelial cells were also in close proximity to Crystallin Alpha B (CRYAB^+^) truncated radial glia (tRG) (**Fig. 1f**, bottom left), a progenitor subtype known to interact with the vasculature during cortical development^37^.

UEA1 endothelial cells were also observed in close association with SOX2 neural progenitors and GFAP astroglial cells (**Fig. 1f**, bottom right). Together, these data show that our high-throughput fusion strategy vascularizes cortical organoids and that the resulting vCAs reconstitute key cellular lineages and re-create salient neurovascular niches of the developing human cortex.

## Single-cell transcriptomic analysis highlights neurovascular cellular diversity in vCAs

To systematically evaluate the cellular composition of our vascularization strategy, we performed single-cell RNA sequencing (scRNA-seq) on VOs, COs, and vCAs generated from three independent hiPSC lines (**Fig. 2a, Extended Data Fig. 2**). After quality controls (Methods, **Extended Data Fig. 2a**), we captured 26,889 high-quality cell transcriptomes (VO, 7,131 cells; CO, 6,521 cells; vCA, 13,237 cells). Transcriptomes were computationally integrated across organoid conditions, clustered, and visualized with uniform manifold approximation and projection (UMAP) plots to permit comparisons of cell composition and lineage across organoids and assembloids (Methods, **Fig. 2b**).

**Fig. 2.**
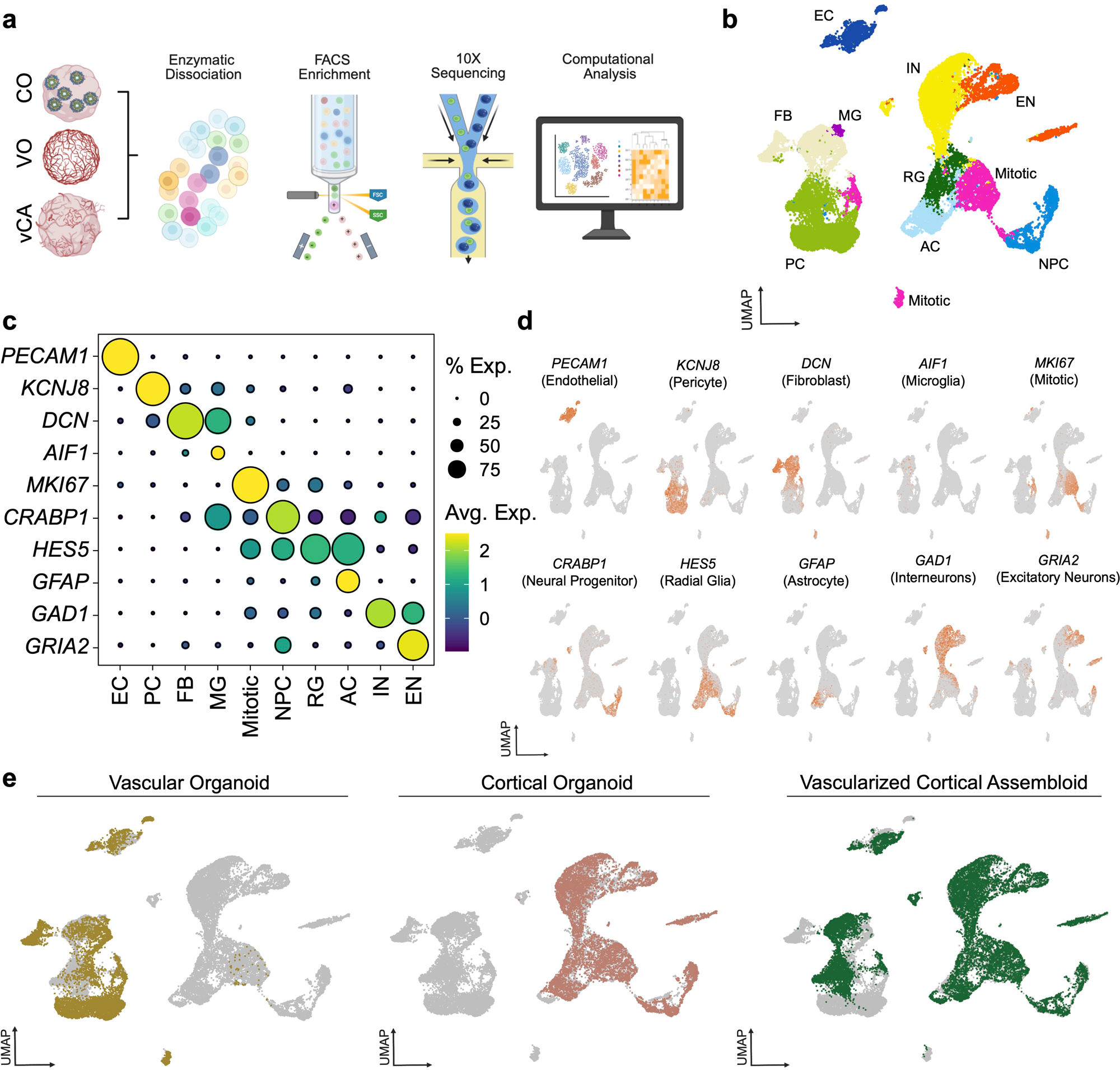
Single-cell transcriptomic profiling demonstrates neurovascular cellular diversity in cortical organoids, vascular organoids, and vascularized cortical assembloids. **a**, Schematic showing experimental workflow for scRNA-seq analysis. Fluorescence activated cell sorting (FACS) was used enrich for CD31^+^ endothelial cells and PDGFRβ^+^ to ensure sufficient vascular capture. **b**, Uniform manifold approximation and projection (UMAP) of 26,889 transcriptomes from vascular organoids (VOs, 7,131 cells), cortical organoids (6,521 cells), and vascularized cortical assembloids (vCAs, 13,237 cells). Three different hiPSC lines were used per condition. UMAP colored by cell identity. EC, endothelial cell; PC, pericyte; FB, perivascular fibroblast; MG, microglia; Mitotic, mitotic cell; NPC, neural progenitor cells; RG, radial glia; AC, astrocyte; IN, interneuron; EN, excitatory neuron. **c**, Dot plot showing expression of cell population marker genes. Avg., average; Exp., expression; Blue, low expression; Yellow, high expression. **d**, Expression distribution of cell population marker genes projected on UMAP embeddings from all transcriptomes as in Gray, low expression; Orange, high expression. **e**, UMAP visualization of all transcriptomes colored by condition. Gold, vascular organoid; Salmon, cortical organoid; Green, vascular cortical assembloids.

Based on canonical marker genes, we identified discrete vascular and neural cell populations, including *PECAM1^+^*endothelial cells, *KCNJ8^+^* pericytes, *DCN^+^* perivascular fibroblasts, *AIF1^+^* microglia cells, *MKI67^+^* mitotic cells, *CRABP1^+^* neuronal progenitor cells (NPCs), *HES5^+^*radial glia cells, *GFAP^+^* astrocytes, *GAD1*^+^ interneurons, and *GRIA2^+^* excitatory neurons (**Fig. 2b-d**). Neural and glial populations predominated in COs and vCAs, whereas vascular and myeloid cells were enriched in VOs and vCAs (**Fig. 2e, Extended Data Fig. 2b**). Each annotated population appeared in organoids and vCAs derived from multiple hiPSC lines (**Extended Data Fig. 2c**), supporting the reproducibility of our differentiation protocols. Importantly, each vCA consistently contained both neural- and vascular-derived populations across all hiPSC lines tested (**Extended Data Fig. 2c**). Thus, vCAs reliably integrate and promote co-development of vascular and neural cell lineages.

### vCAs acquire BBB properties and arteriovenous specification

The BBB is a specialized feature of brain endothelial cells that governs solute and cellular exchange with the circulating blood to support neuronal function ^38,39^. To test whether vascular-cortical integration promotes the development of BBB properties *in vitro*, we compared relevant gene signatures in scRNA-seq datasets from VOs, COs and vCAs (**Fig. 2**). Using “Establishment of BBB” gene ontology (GO:0060856), gene-set scoring revealed higher signature scores in vCAs relative to both organoids (*p* < 0.001), with enrichment in neural- and vascular-derived cell populations (**Extended Data Fig. 3a**). To define the cell-specific basis for BBB development in vCAs, we queried gene sets for “Establishment of BBB” (GO:0060856), and “Maintenance of BBB” (GO:0035633) (**Fig. 3a**). This analysis showed enrichment of select tight junction genes, e.g., *OCLN* (encodes occludin), and critical regulators of endothelial BBB formation, e.g., *MFSD2A* (**Fig. 3a**)^39,40^. Tight junctions are a cardinal feature of the BBB, limiting non-specific paracellular transport between adjacent endothelial cells^39,41^. Quantitative immunostaining confirmed that tight junction proteins, such as zonula occludens-1 (ZO-1) and claudin-5 (CLDN5), were significantly increased in vCAs compared with VOs (*p* <0.001) (**Fig. 3b**). Transmission electron microscopy (TEM) similarly revealed tight-junction complexes in day-95 vCA endothelium (**Fig. 3c**). Endothelial tubes were lumenized and displayed low rates of pinocytosis and vesicular transport (**Fig. 3c**), consistent with reduced transcytosis in the physiologic BBB^14,42-46^. Together, these data support acquisition of organotypic specialization of endothelial cells within vCAs.

**Fig. 3.**
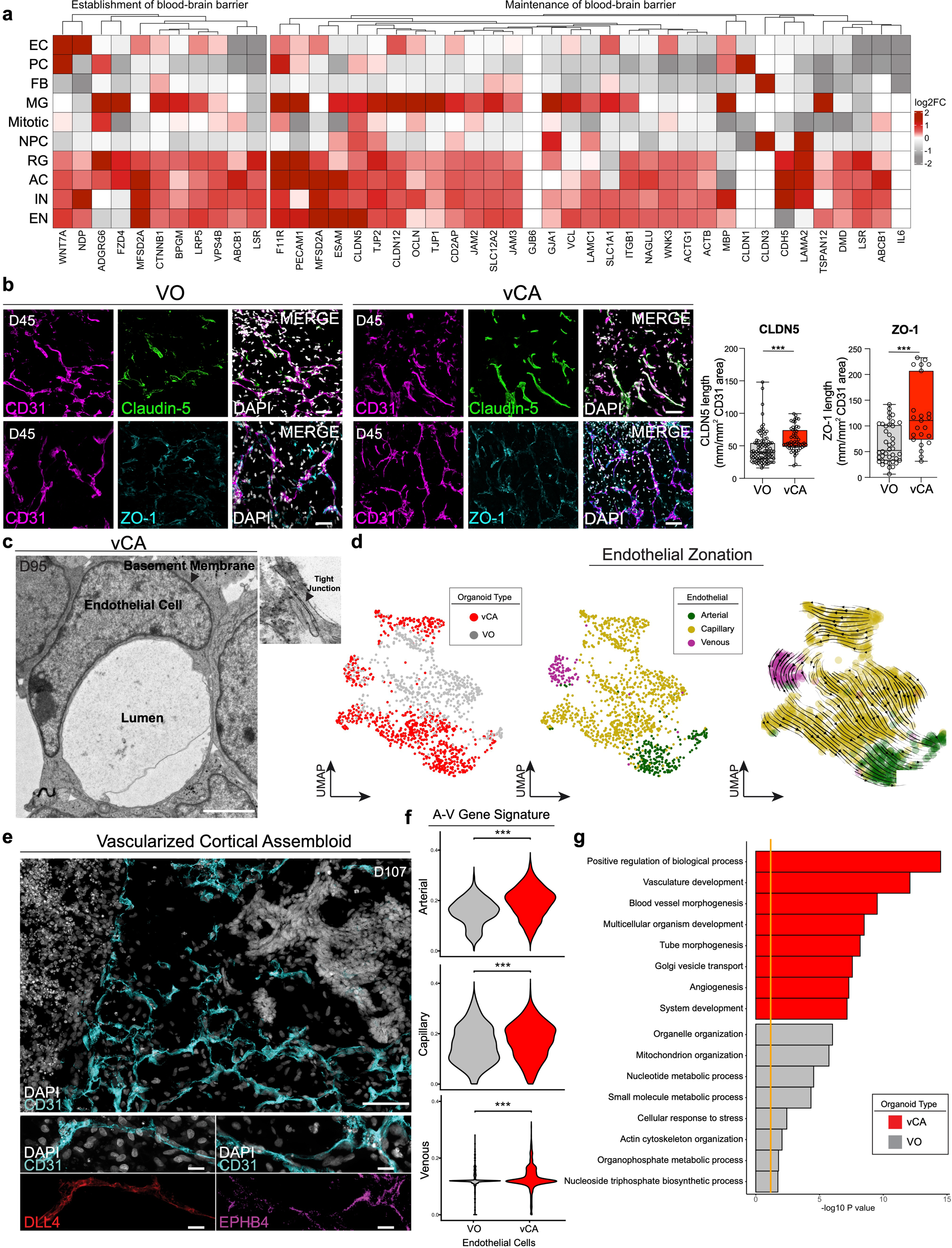
Blood-brain barrier acquisition and endothelial specialization in vascular cortical assembloids. **a**, Heatmap visualization of cell-specific enrichment of establishment and maintenance of blood-brain barrier genes in vascular cortical assembloids (vCAs). Gray, low expression; Red, high expression. EC, endothelial cell; PC, pericyte; FB, perivascular fibroblast; MG, microglia; Mitotic, mitotic cell; NPC, neural progenitor cells; RG, radial glia; AC, astrocyte; IN, interneuron; EN, excitatory neuron. **b**, Representative confocal microscopy analysis showing Claudin-5 (green) and ZO-1 (cyan) tight junctions in CD31^+^ endothelial cells (magenta) in vascular organoids (left, VO) and vCAs (middle). Scale bar, 50 μm. Right, Quantification of endothelial claudin-5 (CLDN5) and ZO-1 tight junction length in VOs and vCAs. n = 3 hiPSC lines per condition, 3-5 organoids per cell line per condition, all replicates shown. ***, *P* < 0.001. **c**, Representative transmission electron microscopy image of day 95 vCA endothelial cell in cross section. Arrowhead shows basement membrane. Inset, high magnification of endothelial tight junction protein complex. Scale bar, 2µm. **d**, UMAP visualization of endothelial cell transcriptomes from VOs and vCAs colored by condition (left), arteriovenous identity (middle), or showing endothelial RNA velocity trajectories (right). Gray, VO; Red, vCA; Green, arterial; Yellow, capillary; Magenta, venous. **e**, Representative confocal microscopy analysis showing DLL4^+^ (arterial, red) and EPHB4^+^ (venous, magenta) endothelial cells in day 107 vCAs. CD31^+^ (cyan) labels all endothelial cells. DAPI (white) labels cell nuclei. Scale bars 100 μm for lower magnification image (top), 10 µm for high magnification images (bottom). **f**, Violin plots for endothelial arterial, capillary and venous gene signatures in VOs (gray) and vCAs (red). ***, *P* < 0.001. **g**, Top gene ontology enrichments for endothelial cells from vCAs (red) and VOs (gray). Orange line, threshold for FDR < 0.05.

During brain development, barriergenesis arises from instructive cues from pericytes and neuroepithelium ^13-15,29,47,48^. In vCAs, the expression of BBB-inducing ligands, e.g., *WNT7a*, were enriched in neuronal subsets and pericytes (*p* < 0.05) (**Fig. 3a**)^13,49,50^. Brain organotypic vascular identity was not limited to endothelial cells, vCA pericytes were also significantly enriched in brain-specific gene signatures compared to VOs (*p* < 0.001), and may support specialized endothelial-pericyte interactions underlying initial barriergenesis (**Extended Data Fig. 3b-c**)^15,51^. Therefore, we next investigated whether BBB maturation signaling cascades were recapitulated in vCAs. Using reciprocal ligand-receptor interactions ^52^, we surveyed cell-to-cell communication pathways in an unbiased manner using scRNA-seq data from vCAs compared to COs and VOs. Pathways over-represented in vCAs included those linked to endothelial barriergenesis (e.g., canonical and non-canonical WNT signaling), pericyte recruitment (e.g., tumor necrosis factor-related apoptosis-inducing ligand (TRAIL) signaling), and arteriovenous specification (e.g., ephrin (EPH-A or -B) signaling) (**Extended Data Fig. 3d-g**)^13,53,54^. Thus, vCAs recapitulate instructive neurovascular cues of human cortical vascular development, and the integrated neurovascular niche is sufficient to induce BBB gene programs and endothelial barriergenesis.

Beyond BBB specialization, we sought to establish whether vCA endothelium is organized according to the arteriovenous hierarchy that is characteristic of the developing cortical vasculature^11,12^. To assess arteriovenous specification, we sorted endothelial cells from VOs and vCAs *in silico*, performed iterative clustering, and plotted curated endothelial arteriovenous gene signatures (**Supplementary Table 3**)^11,12,55,56^. Capillary identity predominated in both models (**Fig. 3d**). Surprisingly, vCAs resolved transcriptionally into distinct arterial and venous endothelial populations that we validated by immunostaining for delta-like protein 4 (DLL4, which denotes arterial CD31^+^ endothelial cells) and EPHB4 (venous endothelial cells) (**Fig. 3d-e**). Arterial, venous, and capillary gene signature scores were significantly enriched in vCAs compared with VOs (*p* <0.001) (**Fig. 3f**), indicating that cortical-derived cues contribute to arteriovenous specification. RNA velocity analyses inferred that endothelial cell trajectories are similar to endothelial specification in primary tissue, beginning as venous endothelial cells, progressing through capillary intermediates, and terminating as arterial endothelial cells^11,57^ (**Fig. 3d**). Differential gene expression analyses showed vCA endothelial cells were enriched in the expression of gene programs underlying vascular development, morphogenesis, and Golgi-vesicle transport (**Fig. 3g**), consistent with vascular maturation. Because arteriovenous specification and maturation coincide with reduced cell proliferation^58,59^, we computationally inferred cell cycle position based on gene expression and observed fewer endothelial cells, pericytes, and perivascular fibroblasts in M/G1 phases in vCAs (*p* < 0.001) (**Extended Data Fig. 3h**)^60^, consistent with quiescence. Collectively, these data show that cortical integration in vCAs promotes BBB specification as well as arteriovenous specialization, maturation, and quiescence, establishing neurovascular features and hierarchical vascular patterning despite the absence of perfusion.

### vCAs exhibit enhanced progenitor organization and neuronal development

Without the vasculature, CO progenitor populations become hypoxic, receive inadequate nutrient delivery, and lose trophic support, which stalls maturation and limits neurodevelopmental fidelity of organoids ^21,22,61,62^. Consistent with prior work ^25,27-29^, quantitative immunostaining for day-104 COs and vCAs revealed reduced hypoxia (HIF-1α^+^cells, *p* < 0.001) and apoptosis (Cleaved Caspase-3^+^ cells, *p* < 0.05) and increased cell proliferation (Ki67^+^ cells, *p* < 0.001) in vCAs (**Fig. 4a**). We next examined progenitor architecture, a hallmark of early corticogenesis^63^. At matched timepoints, immunostaining of SOX2^+^ NPCs showed well-organized and persistent radial rosettes in vCAs, in contrast to more collapsed and disorganized rosettes in COs (**Fig. 4b**). Quantification confirmed larger rosette size and higher cell density in vCAs (*p* < 0.05) (**Fig. 4c**). Thus, vascularization prolongs progenitor viability and preserves progenitor niche architecture, promoting healthier and more advanced/robust neurodevelopment.

**Fig. 4.**
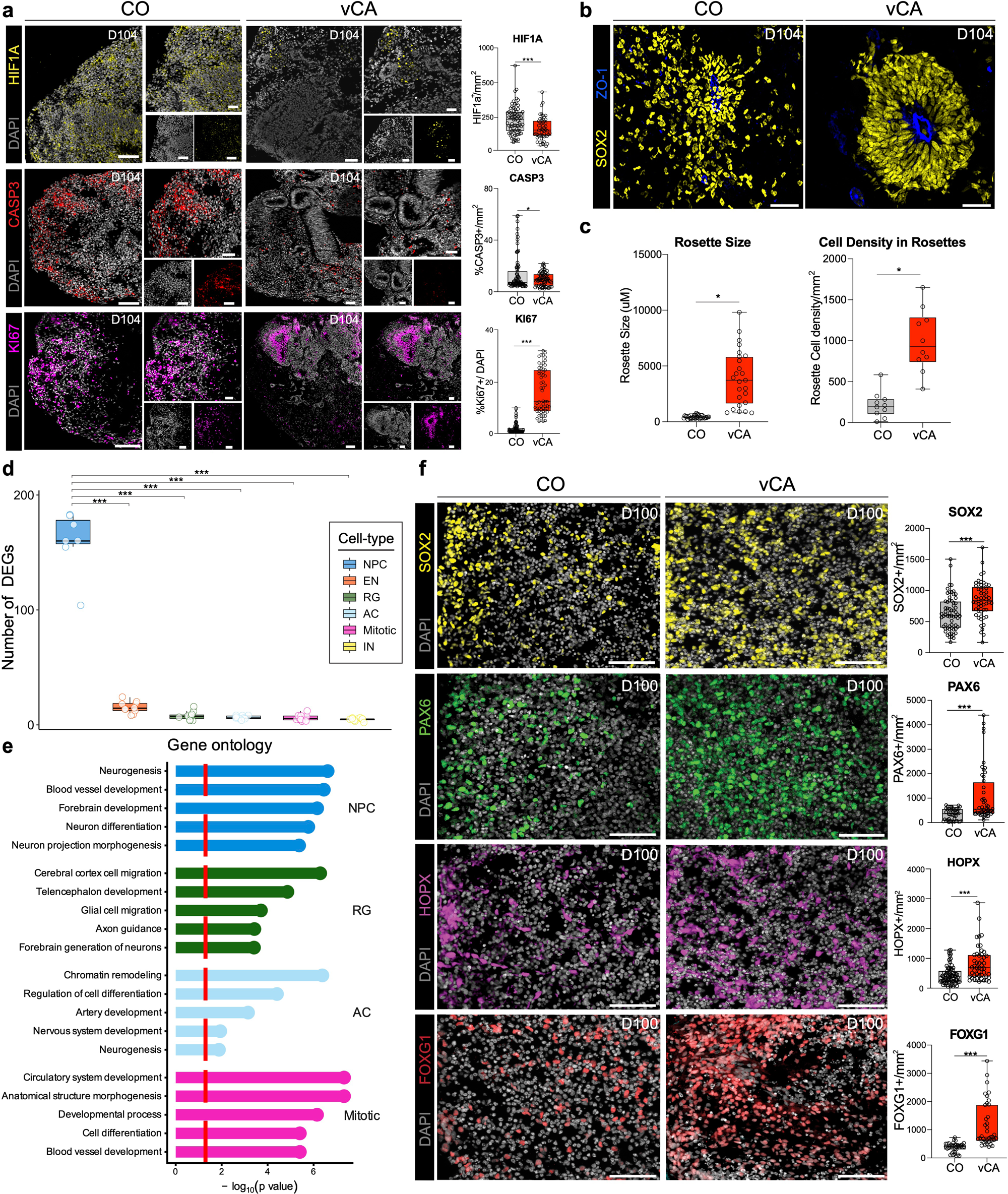
Improved neural progenitor microenvironment and organization in vascularized cortical assembloids. **a**, Representative confocal microscopy analysis showing HIF-1α^+^hypoxic cells (top, yellow), cleaved caspase 3^+^ (CASP3) apoptotic cells (middle, red), and Ki67^+^ proliferative cells (bottom, purple) in day 104 cortical organoids (COs) and vascularized cortical assemblies (vCAs). CD31^+^ (cyan) labels endothelial cells. DAPI (white) labels cell nuclei. Scale bars 100 μm for lower magnification image (left), 50 µm for high magnification images (right). Right, quantification of HIF-1α^+^ hypoxic cells, cleaved caspase 3^+^ (CASP3) apoptotic cells, and Ki67^+^ proliferative cells in COs and vCAs. n = 3 hiPSC lines per condition, 3-5 organoids per cell line per condition, all replicates shown. *, *P* < 0.05; ***, *P* < 0.001. **b**, Representative confocal microscopy analysis showing SOX2^+^ neural progenitor cell (yellow) rosette architecture in day 62 COs and vCAs. ZO-1 (blue) labels the apical surface of the neuroepithelial cells. Scale bars, 50 µm. **c**, Quantification of rosette size (left) and cell density (right) in COs and vCAs. n = 3 hiPSC lines per condition, 3-5 organoids per cell line per condition, all replicates shown. *, *P* < 0.05. **d**, Gene burden analysis of progenitor and glia cell populations between vCAs and COs. ***, *P* < 0.001 compared to all other cell populations (one-way ANOVA with Tukey’s multiple comparisons test). Mitotic, mitotic cell; NPC, neural progenitor cells; RG, radial glia; AC, astrocyte; IN, interneuron; EN, excitatory neuron. **e**, Top gene ontology enrichments from differentially expressed genes in neural progenitor cells (NPC), radial glia (RG), astrocytes (AC), and mitotic cells in vCAs. Dashed red line, threshold for FDR < 0.05. **f**, Representative confocal microscopy analysis showing SOX2^+^ neural progenitor cells (yellow), PAX6^+^ neural progenitor cells (green), HOPX^+^ outer radial glia (magenta), and FOXG1^+^ telencephalic progenitor cells (Red) in day 100 COs and vCAs. Right, quantification of each cell population from immunostaining. n = 3 hiPSC lines per condition, 3-5 organoids per cell line per condition, all replicates shown. ***, *P* < 0.001. Scale bars, 100 µm.

Beyond metabolic support, the vasculature provides trophic support and paracrine cues to promote neurogenesis and cortical development ^4,5,64^. Our scRNA-seq analyses of vCAs compared to VOs and COs revealed that vascular cell lineages expressed genes linked to pro-neurodevelopmental pathways, including telencephalon development, neuronal migration, and axonogenesis (**Extended Data Fig. 4a**). Notably, the strongest enrichment was observed in pericytes (**Extended Data Fig. 4a**), underscoring the importance of incorporating multiple vascular cell types. Within neurons and glia, differential gene expression analysis showed the greatest transcriptional shifts in NPCs (**Fig. 4d, Extended Data Fig. 4b**), supporting an early influence of the vasculature on progenitors in vCAs. Gene ontology analysis of NPC differentially expressed genes (DEGs) showed enrichment for neurogenesis, neuronal differentiation, and morphogenesis, and telencephalic development (**Fig. 4e**). Consistent with the expression of these gene signatures, quantitative immunostaining showed increased abundance in SOX2^+^ and PAX6^+^ NPCs, HOPX^+^ outer radial glia, and Forkhead box G1 (FOXG1^+^) telencephalic progenitors (*p* <0.001, **Fig. 4f**), supporting strengthened cortical identity in vCAs compared with COs. We also observed greater distribution in DCX^+^ neuroblasts in vCAs, supporting improved neurogenesis in the assembloid model **(Extended Data Fig. 4c)**. Together, these data show that vascularization creates a pro-neurogenic niche that expands and organizes cortical NPCs and strengthens cortical identity, improving neurodevelopmental fidelity in vitro.

### Vascularization promotes neuronal maturation and functional connectivity in vCAs

Because organoid vascularization that is achieved through xenotransplantation improves neuronal maturation, longevity, and axonal outgrowth^23,24^, we next tested whether vascularization similarly promotes neuronal maturation and connectivity *in vitro*. scRNA-seq differential gene expression analysis showed that vCA excitatory neurons and interneurons upregulated the expression of maturation gene programs, including neuron projection morphogenesis (excitatory), neuron differentiation (excitatory), axon development (interneurons), and synapse organization and assembly (interneurons), relative to COs (**Fig. 5a**). To examine whether these transcriptional shifts are associated with morphological correlates, we performed quantitative immunostaining in day 104 vCAs and COs. Compared to Cos, vCAs contained significantly greater microtubule-associated protein 2 (MAP2^+^) (*p* < 0.05) and NeuN^+^ mature neurons than COs (*p* < 0.001) (**Fig. 5b**). vCAs also showed increased abundance of SATB2^+^ neurons, which were positioned further from apical rosettes (*p* < 0.001) (**Fig. 5b**), suggesting the emergence of upper-layer neurons and a more advanced cortical developmental trajectory. On average, vCA neurons exhibited significantly larger somata (1.2-fold, *p* < 0.001) and greater dendritic and axonal neurite density, as quantified by MAP2 (dendrites) and neurofilament medium polypeptide (NFM; axons) expression (*p* < 0.001) (**Fig. 5c-d**), consistent with more elaborate arborization. Pre- and post-synaptic staining for synaptophysin (SYP) and HOMER1, respectively, showed higher synapse density in vCAs than COs (**Figure 5d, Extended Data Fig. 4d**). Therefore, vCA neurons exhibit transcriptional and morphological changes consistent with enhanced maturation and synaptogenesis compared to COs.

**Fig. 5.**
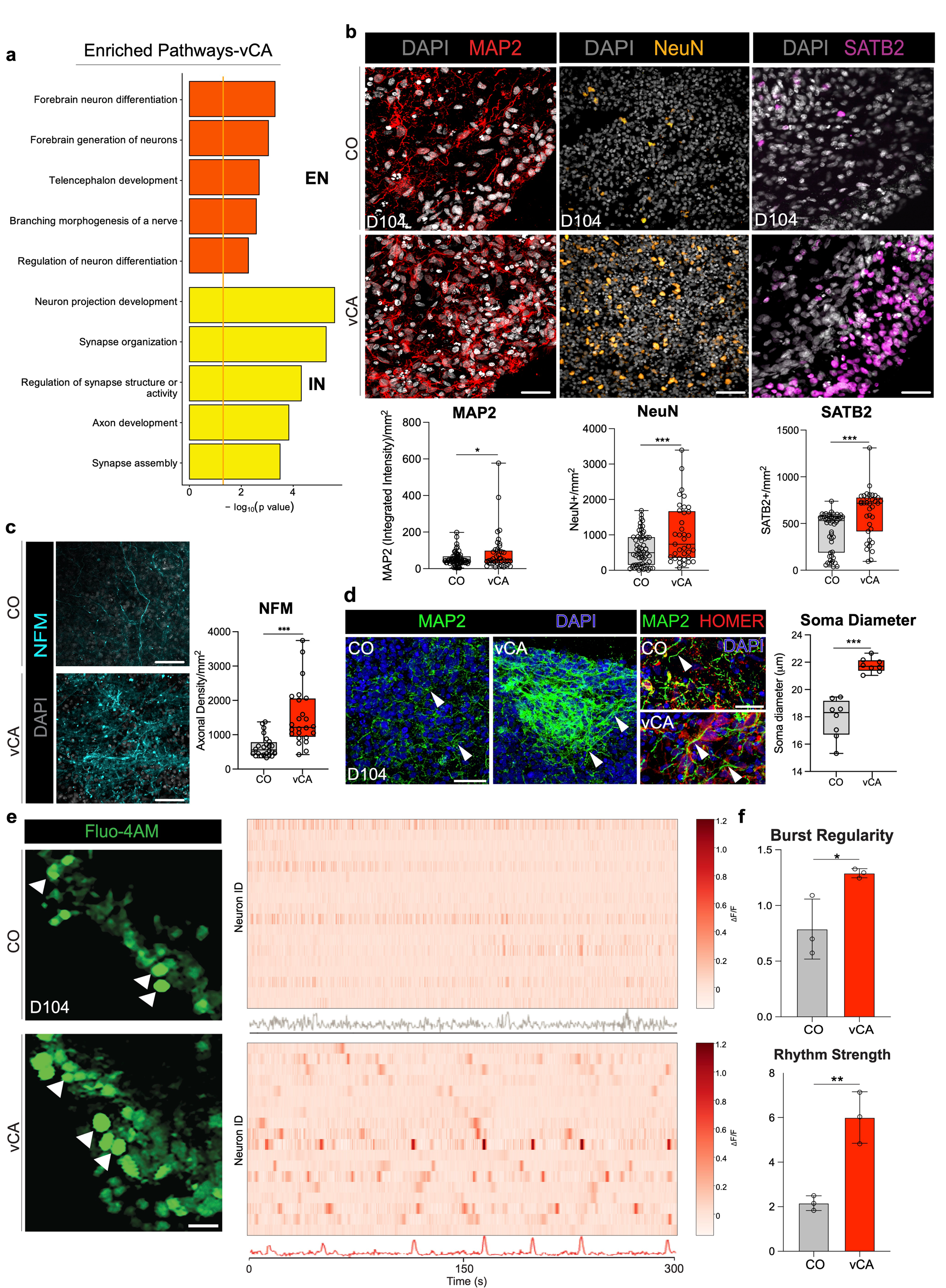
Neuronal maturation and functional connectivity in vascularized cortical assembloids. **a**, Top gene ontology enrichments from differentially expressed genes in excitatory neurons (EN) and interneurons (IN) in vascularized cortical assembloids (vCAs). Orange line, threshold for FDR < 0.05. **b**, Representative confocal microscopy analysis showing neuronal maturation markers MAP2 (red) and NeuN (orange) and SATB2^+^ upper-layer-like neurons (magenta) in day 104 cortical organoids (COs) and vascularized cortical assembloids (vCAs). DAPI (white) labels cell nuclei. Scale bars, 50 µm. Bottom, quantifications of MAP2^+^ integrated density (left), NeuN^+^ cell counts (middle), and SATB2^+^ neurons (right) in COs (gray) and vCAs (red). n = 3 hiPSC lines per condition, 3-5 organoids per cell line per condition, all replicates shown. *, *P* < 0.05; ***, *P* < 0.001. **c**, Representative confocal microscopy analysis showing NFM^+^ axons in day 104 COs and vCAs. Right, quantification of NFM^+^ axonal density in COs (gray) and vCAs (red). n = 3 hiPSC lines per condition, 3-5 organoids per cell line per condition, all replicates shown. *, *P* < 0.05; ***, *P* < 0.001. Scale bars, 50 µm. **d**, Representative confocal microscopy showing MAP2^+^ neuronal dendrites and soma (green) and HOMER^+^ post-synaptic densities (Red) in COs and vCAs. DAPI (blue) labels cell nuclei. Arrowheads in lower magnification images, MAP2^+^ dendrites. Arrowheads in higher magnification images, co-localization. Scale bars, 50 µm. **e**, Representative microscopy of Fluo-4AM functional calcium imaging in COs and vCAs. Arrowheads, active neurons. Right, temporal raster plots and aggregate field potentials of spontaneous calcium transients in COs (top) and vCAs (bottom). Scale bars, 20 µm. **f**, Quantification of burst regularity and rhythm strength of calcium transients in COs (gray) and vCAs (red). n = 3 organoids per condition. *, *P <*0.05; *\*\*, P <* 0.01.

This improved neurodevelopment prompted us to investigate patterns of neuronal activity in vCAs. Functional calcium imaging revealed greater consistency of spontaneous calcium transients in vCAs, whereas COs exhibited prolonged burst-suppression intervals (**Fig. 5e**). Using established quantitative network-level analyses^65^, vCAs displayed increased burst regularity (p < 0.05) and rhythm strength (*p* < 0.01) (**Fig. 5f**), indicating greater temporal coordination of neuronal discharges, consistent with mature cortical networks. Together, these data show that vascularization improves neuronal maturation and circuit assembly, thereby improving neuronal function and connectivity in cortical assembloids.

### vCAs exhibit improved fidelity to human developing cortical tissue compared to other organoid models

The fidelity of existing cortical organoid protocols to primary human tissues remains debated^22,66,67^. To compare vCA fidelity with the developing human cortex, we integrated our scRNA-seq data with two developmental neurovascular atlases using scVI^11,12,68,69^(**Fig. 6a**). The integrated UMAP showed that organoid-derived cells broadly aligned with canonical primary brain cell lineages (**Fig. 6b-c, Extended Data Fig. 5a**), supporting conservation of vascular and neural specification programs in vCAs. When visualized by source, excitatory neurons and interneurons from vCAs and primary tissues were closely co-localized (**Fig. 6c**), supporting physiological maturation and appropriate cortical trajectories. Marker gene expression identified cell identities across tissue sources, including *PECAM1^+^*endothelial cells, *KCNJ8^+^* pericytes, *DCN^+^* perivascular fibroblasts, *AIF1^+^* microglia, *MKI67^+^* mitotic cells, *CRABP1^+^* NPCs, *HES5^+^*radial glia, *GFAP^+^* astrocytes, *GAD1*^+^ interneurons, *GRIA2^+^* excitatory neurons, and *MBP^+^*oligodendrocytes (**Fig. 6d**), confirming broad lineage fidelity with primary human cortical tissue and a complex neurovascular niche in vCAs.

**Fig. 6.**
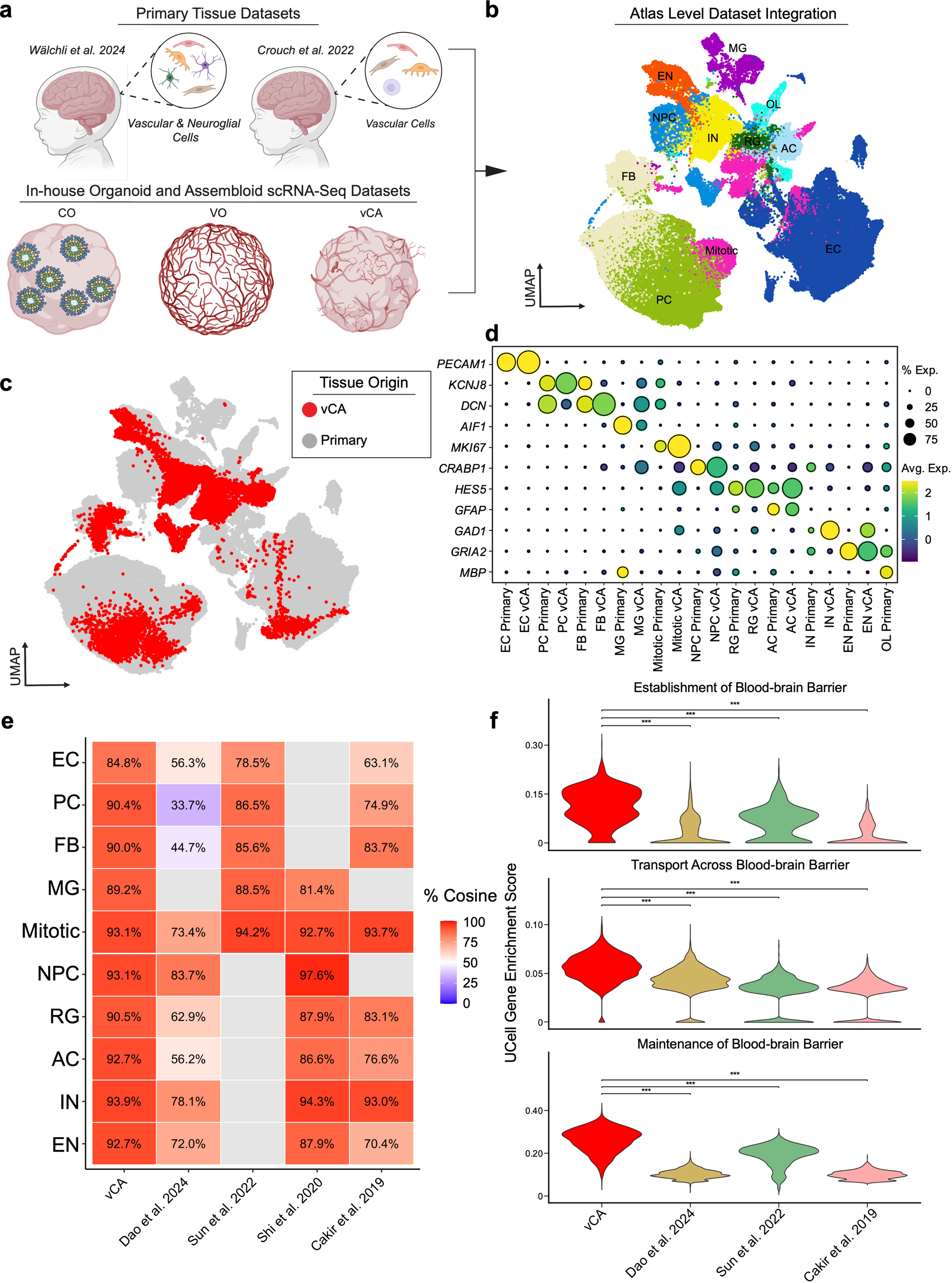
Fidelity and cross-protocol benchmarking of vascularized cortical assembloids. **a,** Schematic illustrating primary tissue datasets used for integration with organoid scRNA-seq datasets. **b-c**, UMAP visualizations of 22 transcriptomes following atlas-integration colored by cell identity (B) or tissue origin (C). EC, endothelial cell; PC, pericyte; FB, perivascular fibroblast; MG, microglia; Mitotic, mitotic cell; NPC, neural progenitor cells; RG, radial glia; AC, astrocyte; IN, interneuron; EN, excitatory neuron. Gray, primary tissue; Red, vascularized cortical assembloid (vCA). **d**, Dot plot showing expression of cell population marker genes in human primary tissues and vCAs. **e**, Heatmap showing cell-specific transcriptional fidelity across published organoid vascularization protocols with available scRNA-seq data. Red, more similar; Blue, less similar. **f**, Violin plots of blood-brain barrier establishment, transport, and maintenance gene sets across published organoid vascularization protocols with available endothelial scRNA-seq data. ***, *P* < 0.001 compared to all other protocols (one-way ANOVA with Tukey’s multiple comparisons test).

We next benchmarked our approach to other published organoid vascularization protocols using established methods ^25,27-29,69-73^(Methods) (**Extended Data Fig. 5b, Supplementary Table 4**). Cell type annotations were harmonized across scRNA-seq datasets from four vascularized organoid models to facilitate cross-study comparisons (**Extended Data Fig. 5b**). We then constructed pseudobulk profiles and scored cell type-specific gene expression from each model against profiles from primary tissue^11,12^. Transcriptional fidelity varied across protocols and cell lineages; however, our in-house vCAs most closely matched primary tissue cell populations and captured the most complete complement of vascular, neural and glial cell types (**Fig. 6e**). Our vCAs performed best in vascular cells (endothelial 84.8%, pericyte 90.4%, fibroblast 90.0%) and glia (astrocyte 92.7%, radial glia: 90.5%) (**Fig. 6e**). Additionally, vCA endothelial cells showed statistically the greatest enrichment for BBB establishment, maintenance, and transport gene signatures relative to the other protocols with available endothelial cell scRNA-seq data (**Fig. 6f**). Thus, vCAs closely recapitulate diverse primary cortical cell states with better cell diversity and advanced neurovascular properties, including enriched BBB programs, outperforming alternative protocols and providing a high-fidelity, high-throughput platform to more faithfully model human cortical neurovascular co-development.

## Discussion

Here, we present a modular assembloid strategy that vascularizes cortical organoids at scale leading to high fidelity neurodevelopment. By generating mesoderm-derived vascular organoids in 96-well format and fusing them to cortical organoids in pre-fabricated molds, we achieve symmetric, reproducible vascular integration that resembles the angioarchitecture of the developing human cortex^28,29^. The vasculature that forms contain lumens and re-capitulates core cerebrovascular properties, including multi-lineage cell composition, BBB features, and arteriovenous specification. Vascularization preserves neuroepithelial architecture, provides trophic support, and reduces hypoxia and metabolic stress, thereby expanding NPC populations and promoting neuronal maturation and circuit connectivity to more closely approximate the developing human cortex.

Our approach advances both the fidelity and throughput of vascularized brain organoid models. Rodent xenotransplantation yields perfused host vasculature and similarly improves neuronal maturation, longevity, and circuit assembly, but this approach limits human specificity in vascular and immune cells, introduces host-graft variables, and constrains scale and vessel-specific disease modeling^23,24^. Cortical organoid-endothelial co-culture strategies, e.g., using HUVECs or ETV2-driven endothelialization, form vessel-like meshes and some BBB features, but they rely on non-brain endothelium or reprogrammed neuroectoderm and underrepresent perivascular cell lineages, reducing physiological relevance ^25,27^. Prior neural-vascular fusion assembloids improve cell representation, but often show asymmetric vascularization, non-physiologic geometry, and remain limited in benchmarking to primary developmental tissue^28,29^. By contrast, our 96-well, mold-guided fusion produces symmetric, reproducible vascular networks that resolve both arterial and venous endothelial cell states in vitro (beyond the venous states previously reported) ^29^. Cell type comparisons to primary human datasets and cross-protocol benchmarking demonstrate our vCAs generate the greatest organotypic concordance across endothelial cells, pericytes, and perivascular fibroblasts, including more robust BBB programs, and show concurrent gains in neuroepithelial organization, progenitor output and network activity, all without xenografting.

## Limitations of the study

The vasculature in our vCAs - and any vascularized organoid model - is not perfused. Without physiological flow, blood vessels cannot dynamically match oxygen or nutrient delivery to organoid metabolic needs, as in the human brain^74^. Shear stress also promotes cerebrovascular maturation, enhancing BBB development and integrity as well as promoting arteriovenous specification^75,76^. While vCAs contain microglia, they lack the broader immune complement present in the human brain^55,77^. Our characterizations were also performed at a single time-point, so whether vascularization enables or accelerates long-term maturation, including postnatal specializations, remains unresolved ^78^. Future work should incorporate bioengineered perfusion to achieve physiological flow modeled on the human brain, expand immune cell complexity, and extend modular neural-vascular fusion to patterned regional organoids, multi-region assembloids, and later developmental stages to probe region-specific neurovascular development, specialization, and maturation.

Despite these limitations, our vCA platform reconstitutes human neurovascular co-development with higher fidelity than other reported models, unifying BBB acquisition, arteriovenous specialization, and a pro-neurogenic niche within a reproducible, high-throughput workflow. Coupling its modular design with single-cell benchmarking and convergent structural and functional phenotyping, it provides a practical platform to dissect mechanisms of human neurovascular development, model BBB and vascular dysfunction in disease, and accelerate therapeutic discovery in developmental neurovascular diseases.

## Methods

### Ethics Statement and Tissue Acquisition

Human induced pluripotent stem cell (hiPSC) lines WTC11, 1323-4, and KOLF2.1 were obtained from the Nowakowski and Kriegstein laboratories at the University of California, San Francisco (UCSF) with protocols approved by the institutional review board. Clinical metadata for each line is detailed in (**Supplementary Table 1)**. Second-trimester human developmental brain tissue samples were acquired following approval by the UCSF Human Gamete, Embryo, and Stem Cell Research Committee (Study Number 10-05113). Written informed consent was obtained prior to the procedure for research use, adhering strictly to legal, institutional, and ethical regulations. All samples were de-identified; sex information was not available. All experiments involving primary tissue were performed in accordance with approved protocol guidelines.

### Cell Line Maintenance

hiPSC lines were expanded using Matrigel® (Corning) coated six-well plates in mTeSR1™ Plus medium (Stem Cell Technologies) supplemented with 10 µM Y-27632 ROCK inhibitor (Tocris) for initial thawing only. Cells were maintained in a sterile incubator at 37°C with 5% CO2. Media changes were performed every other day. Cultures were passaged upon reaching approximately 70% confluency using ReLeSR™ (Stem Cell Technologies) combined with manual lifting. Residual cells were plated onto freshly coated plates or cryopreserved by freezing in mFreSR™ (Stem Cell Technologies) at -80°C for 24 hours before transferring to liquid nitrogen for long-term storage. All iPSC lines utilized in this study were between passage 11 and 26.

### Generation of GFP-labeled hiPSCs

hiPSC-GFP cell lines were generated using lentiviral transduction. Second-generation lentivirus was produced by transfecting HEK293T cells cultured at 80-90% confluency in DMEM (Stem Cell Technologies) supplemented with 10% FBS (Fisher Scientific) and 1% Penicillin-Streptomycin (ThermoFisher). Transfection utilized the transfer vector pFUGW (Addgene plasmid #14883), the packaging plasmid pCMV-dR8.2 dvpr (Addgene plasmid #8455), and the envelope plasmid pMD2.G (Addgene plasmid #12259) using nucleofection (FuGENE® HD Transfection Reagent, Promega). HEK293T cells were maintained at 37°C with 5% CO2. Supernatant containing viral particles was collected at 24- and 48-hours post-transfection, pooled, and filtered through a 0.45 µm filter (FischerScientific). Virus-containing supernatant was concentrated by ultracentrifugation at 26,000 × g for 2 hours at 4°C. The viral pellet was resuspended in sterile PBS (Fisher Scientific), aliquoted and stored at -80°C. hiPSCs BU3 & KOLF2.1 were subsequently transduced with the concentrated virus.

### Creation of Cortical Organoid Cell Suspensions

Brain organoids exhibiting forebrain regionalization were generated using a modified differentiation protocol (Pasca et al. 2015). hiPSC lines WTC11, 1323-4, and KOLF2.1 were dissociated into single cells using Accutase (Sigma Aldrich). Following dissociation, cells were resuspended in neural induction medium (NIM) at a density of 12,000 cells per well in 96-well V-bottom ultra-low adhesion plates (Sbio) on day 0 (D0). NIM consisted of GMEM (Fischer Scientific) supplemented with 20% Knockout™ Serum Replacer (KSR, Gibco), 1X MEM Non-Essential Amino Acids Solution (NEAA, Fischer Scientific), 0.11 mg/mL Sodium Pyruvate (Fischer Scientific), 1X Penicillin-Streptomycin (Fischer Scientific), 0.1 mM β-Mercaptoethanol (Fischer Scientific), 5 µM SB431542 (Tocris), 100 nM LDN-193189 (Sigma Aldrich), and 3 µM IWR1-endo (Cayman Chemicals). The medium was renewed every 3 days. Cells were treated with 20 µM Y-27632 ROCK inhibitor for the first 6 days of culture. On D9, organoids were transitioned into maintenance medium 1 (MM1) consisting of a 1:1 mixture of DMEM/F12 (Stem Cell Technologies) containing 1X GlutaMAX™ Supplement (Thermo Fisher Scientific) and Neurobasal™ Medium (Fisher Scientific), supplemented with 2% B-27™ Supplement without Vitamin A (Fisher Scientific), 1% N-2 Supplement (Thermo Fisher Scientific), 1% NEAA, 0.09% of 55 mM β-Mercaptoethanol stock, and 1% Antibiotic-Antimycotic (Fisher Scientific). MM1 was supplemented with 10 ng/mL FGF basic (FGF-2, R&D Systems) and 10 ng/mL EGF (R&D Systems). On D18, organoids were transferred to ultra-low adhesion 6-well plates (Corning) and placed on an orbital shaker set to 90 RPM. Organoids were maintained in MM1 with FGF and EGF supplementation renewed every 3 days until D25. On D35, organoids were transitioned to maintenance medium 2 (MM2), which consisted of a 1:1 mixture of DMEM/F12 with GlutaMAX™ and Neurobasal™ medium, supplemented with 2% B-27™ Supplement with Vitamin A (Thermo Fisher Scientific), 1% N-2 Supplement, 1% NEAA, 0.09% of 55 mM β-Mercaptoethanol stock, and 1% Antibiotic-Antimycotic. MM2 was supplemented with 1:100 dilution of 20 mM L-Ascorbic acid stock (Final concentration 200 µM, Millipore Sigma). Media was changed every three days. Recombinant human LIF protein (10 ng/mL, Alomone Labs) was added to the culture medium between approximately D56-D63. Between weeks 9 and 10 (approximately D63-D70), LIF was replaced with 10 ng/mL BDNF (Alomone Labs) and 10 ng/mL NT-3 (Alomone Labs). From D70 onwards, organoids were maintained in complete MM2 solely supplemented with ascorbic acid.

### Creation of Vascular Organoid Cell Suspensions

Vascular organoids (VOs) were generated using GFP-labeled human induced pluripotent stem cells (hiPSCs) following a modified version of a previously established protocol^31^. Briefly, hiPSCs were maintained on Matrigel®-coated 6-well plates in mTeSR1™ Plus medium within a sterile incubator at 37°C and 5% CO2. On day -1, hiPSC colonies were dissociated into single cells using Accutase. Cells were resuspended at a density of approximately 1.2 × 10 cells per 200 µL of mTeSR1™ Plus medium supplemented with 10 µM Y-27632 ROCK inhibitor. This cell suspension was plated into 96-well V-bottom ultra-low attachment plates to facilitate the formation of embryoid bodies (EBs). On D0, EBs were treated with mesoderm induction medium for three days. This medium consisted of an N2B27 medium base supplemented with 12 µM CHIR99021 (Tocris Biosciences) and 2 ng/mL BMP-4 (Miltenyi Biotec). The N2B27 medium base was composed of a 1:1 mixture of DMEM/F12 and Neurobasal™ Medium (Thermo Fisher Scientific), supplemented with 2% (v/v) B-27™ Supplement, 1% (v/v) N-2 Supplement (Thermo Fisher Scientific), 0.5% (v/v) GlutaMAX™ Supplement, 35 µL of 55 mM β-Mercaptoethanol and 1% (v/v) Penicillin–Streptomycin. The N2B27 base medium was prepared fresh and stored at 4°C for a maximum of two weeks prior to use. Y-27632 was omitted from the mesoderm induction medium. On D3, the medium was replaced with a vascular induction medium, which consisted of the N2B27 medium base supplemented with 100 ng/mL VEGF-A (Peprotech) and 2 µM Forskolin (Sigma Aldrich). Cultures were maintained in this medium until D6, leading to the formation of vascular spheroids (VS). On D6, individual VSs were collected and embedded within 50 µL droplets of cooled Matrigel®. These VO-Matrigel® droplets were incubated at 37°C for 1 hour to permit gel solidification. Solidified droplets were subsequently transferred into 6-well ultra-low attachment plates. 3 mL of pre-warmed (37°C) vascular organoid medium were added to each well. This medium contained StemPro™-34 SFM (Thermo Fisher Scientific) supplemented with 100 ng/mL VEGF-A, 100 ng/mL FGF-2, and 15% FBS. The medium was renewed every two days. VOs were maintained on an orbital shaker at 90 RPM. Blood vessel network formation and sprouting were typically observed starting from D10.

### Fusion of Vascularized Cortical Assembloids

To generate vascularized cortical assembloids (vCAs), D16 VOs were co-cultured with D70 COs. Two VOs were assembled with one CO (2:1 ratio) and embedded together using organoid embedding sheets (Stemcell Technologies, #08579) within a 50 µL Matrigel® droplet and incubated at 37°C for 1 hour. Subsequently, the embeded organoids were removed from the embedding sheet using media (drop-wise) and placed in an ultra-low attachment 24-well plate (Corning). vCAs were cultured in 1:1 StemPro™:MM2 medium (formulation described above), supplemented with VEGF, FGF at 100 ng/mL final concentration, respectively and 1:100 dilution of 20 mM L-Ascorbic acid at 37°C with 5% CO2. Plates were maintained on an orbital shaker at 90 RPM to promote nutrient exchange.

### Immunofluorescent Staining

VOs, COs and vCAs were processed for immunohistochemistry by washing with 1X Phosphate-Buffered Saline (PBS) (Fisher Scientific) and subsequently fixed in 4% paraformaldehyde (PFA) (Sigma-Aldrich) for 30 minutes at room temperature. Following fixation, organoids were washed three times with 1X PBS using a rocker to ensure removal of PFA. For cryopreservation, organoids were submerged in 30% sucrose (Sigma Aldrich) and incubated overnight at 4°C. Organoids were then rinsed in a solution comprising three parts 30% sucrose and two parts Optimal Cutting Temperature (O.C.T) compound (Fisher Scientific), embedded in O.C.T. compound, and sectioned onto charged slides (Fisher Scientific) at a thickness of 15 μm. Slides were dehydrated at 50°C for 10 minutes and allowed to cool to room temperature for 10 minutes. Antigen retrieval was performed by incubating slides with a 1:50 dilution of boiling citrate-based antigen retrieval buffer (pH 6.0, ThermoFisher) for 15 minutes at room temperature. Slides were then washed three times with 1X PBS for 15 minutes each at room temperature. Non-specific binding was blocked by incubating sections with a donkey blocking buffer (DBB) containing 10% donkey serum (Fisher Scientific) in 1X PBS for 1 hour at room temperature. Primary antibodies were diluted 1:250 in DBB and applied to the sections, followed by overnight incubation at 4°C in a light-protected humidified chamber. Sections were washed three times for 5 minutes each with 1X PBS containing 0.1% Triton-X 100 (Fisher Scientific) to remove unbound primary antibodies. Secondary antibodies and DAPI (Thermo Scientific) were diluted 1:500 and 1:1000, respectively, in DBB, applied to sections, and incubated for 2 hours at room temperature in the dark. Following incubation, slides were washed three times for 5 minutes each with 1X PBS. Coverslips were mounted using VectaMount Express Mounting Medium (Vector Laboratories). Slides were allowed to dry at room temperature for 20 minutes, stored overnight at 4°C, and imaged the following day using a NIKON Eclipse Ti2 microscope.

### Calcium Imaging and Functional Assessment

To depolymerize the extracellular matrix in the vascularized cortical assembloids, organoids were transferred into a cell recovery solution (Corning) and incubated for 20 minutes at 4°C. Both vascularized cortical assembloids and brain organoids were washed with PBS (Thermo Fisher) three times for 5 minutes. 40 µL of Fluo-4AM dye (Fisher Scientific) and 16 µL of Pluronic F-127 (Thermo Scientific) were added to 8 mL of Hanks Balanced Salt Solution (Sigma Aldrich). This solution was added to the organoids for 1 hour at 37°C. Following incubation, the organoids were washed with Hanks Balanced Salt Solution three times for 5 minutes. The organoids were then added back into their conditioned media and transported for imaging. 5 minute videos of neuronal firing were obtained using the Nikon A1R LSM and MP microscope.

### Transmission Electron Microscopy

To functionally validate the neurovascular unit within our vascularized cortical assembloids, we performed transmission electron microscopy on D95 vascularized cortical assembloids. Organoids were removed from culture, washed with PBS (Thermo Fisher) and transferred to 2% glutaraldehyde in 0.1 M sodium cacodylate buffer (pH 7.2-7.4), wrapped in parafilm and stored at 4°C overnight. Organoids were rinsed with 0.1 M sodium cacodylate buffer three times for 5 minutes each. The organoids were incubated in 1% Osmium tetroxide + 1.6% Potassium ferricyanide (KFECn) in 0.1 M sodium cacodylate buffer. The organoids were again rinsed three times with 0.1 M sodium cacodylate buffer for 5 minutes each, followed by a wash with distilled water for 5 minutes. Organoids were then incubated in 0.5% Uranyl acetate for 1 hour in the dark. Following rinse with distilled water three times for 5 minutes each, organoids were dehydrated with ascending concentrations of acetone for 10 minutes each (35%, 50%, 75%, 80%, 95%, 100%, 100%). Incubations in a 2:1 mixture of acetone and resin with accelerator (BDMA), respectively, followed for 1 hour. Organoids were then incubated in a 1:1 mixture of acetone and resin with accelerator, respectively, for 1 hour. This followed an additional incubation using 1:2 mixture of acetone and resin with accelerator, respectively, overnight. Organoids were incubated in 100% resin with accelerator three times for 2 hours each. The samples were embedded into appropriate molds and baked in an oven at 60°C for 2 days. Samples were sectioned using the Leica EM UC7 microtome and subsequently imaged on the FEI TECNAI 12 transmission electron microscope.

### Confocal Image Analysis

For quantitative imaging experiments, images of entire tissue sections were acquired as z-stacks (8-10 μm thickness, ensuring comprehensive tissue coverage) using NIKON Eclipse Ti2 series confocal microscope. Maximum intensity projection images were subsequently reconstructed from these z-stacks for analysis. Quantification was performed on four representative, randomly selected fields from three non-adjacent tissue sections per sample. All analyses derived from at least 3 independently generated organoid structures for each experimental condition. Cell counts obtained from each field were normalized to the corresponding imaged tissue area (μm²), typically determined using CellProfiler. Intensity quantifications were conducted by utilizing the "MeasureObjectIntensity" module in CellProfiler (v.4.2.8). A student t-test was used to compare quantitative evaluation using GraphPad Prism software v.10.4.2. P-values < 0.05 were considered statistically significant. Outliers were excluded based on being above or below 1.5 times the interquartile range.

### Quantitative Analysis of Vascular Network Morphology

To quantitatively assess vascular network characteristics, immunofluorescence images of vCAs and primary fetal brain tissue sections, stained for the endothelial marker CD31, were analyzed using the AngioTool software package (Zudaire et al. 2011). Input images were processed through the AngioTool analysis pipeline, which performs vessel identification utilizing multiscale Hessian-based filtering, followed by segmentation and skeletonization of the vascular structures. For each analyzed image, key morphometric parameters were computed by the software, including total vessel length (mm), vascular density (calculated as the percentage of total area occupied by vessels), and branching index (defined as the number of vessel junctions per mm²). Vessel identification and segmentation parameters within AngioTool were carefully optimized for each dataset to ensure accurate representation of the vascular networks prior to automated quantification. The derived quantitative data for total vessel length, vascular density, and branching index were subsequently used for comparative statistical analysis between experimental groups.

### Single-cell Dissociation and FACS Enrichment

Organoids were dissociated into single-cell suspensions using a modified version of a previously described protocol ^3^. Briefly, 7-8 organoids per condition were collected, washed with DPBS, and mechanically dissociated into approximately 1 mm² pieces. Tissue fragments were incubated in a collagenase/dispase solution (3 mg/mL, Thermofisher) for 30 minutes at 37°C. Samples were further triturated in Hank’s Balanced Salt Solution (HBSS) buffer (Sigma Aldrich), supplemented with 1% Bovine Serum Albumin (BSA), 0.1% glucose (Sigma Aldrich) containing 0.25 mg/mL DNase I (Roche) for 30 minutes at 37°C. The resulting cell suspension was filtered through a 70 μm cell strainer (Fisher Scientific) and centrifuged at 200 × g for 5 minutes at 4°C. Cell pellets were resuspended in Stain Buffer (0.2% BSA with 0.09% Sodium Azide, Sigma Aldrich). To enrich for vascular cell populations, cells were incubated with FITC-conjugated anti-CD31 (1:100 dilution, BD Pharmingen, cat. no. 555445) and PE-conjugated anti-CD140b (PDGFR-β) (1:200 dilution, BD Pharmingen, cat. no. 558821) antibodies for 30 minutes on ice. DAPI (1:400 dilution, Invitrogen) was added for live/dead cell discrimination. Fluorescence-Activated Cell Sorting (FACS) was performed using a BD FACS Aria II Flow Sorter (BD Biosciences). Data analysis was conducted using FlowJo software v10.10.

### Single-cell RNA Sequencing

Single-cell RNA sequencing (scRNA-seq) libraries were prepared using the Chromium Next GEM Single Cell 3’ Reagent Kits v3.1 (10x Genomics Inc.). Enriched vascular cell populations were sorted directly into a sequencing buffer (DPBS containing 10% Fetal Bovine Serum (FBS) and 1% BSA) at a target concentration of 1000 cells/μL, aiming for a recovery of approximately 10,000 cells per sample. Library generation followed the manufacturer’s recommended protocol. Briefly, single cells were partitioned into Gel Bead-In-Emulsions (GEMs) for barcoding, followed by reverse transcription, cDNA amplification, and sequencing library construction. Libraries were pooled and sequenced on an Illumina NovaSeq X platform (Illumina Inc.) to an approximate depth of 4.2 × 10 reads per sample.

### Single-cell RNA Sequencing Data Processing

Raw sequencing data (FASTQ files) from three distinct organoid types (cortical organoid, vascular organoid, and vascularized cortical assembloids) generated across two independent experiments were processed using Cell Ranger (v8.0.1, 10x Genomics Inc.). Downstream analysis was performed using the Seurat package (v5.1) in R (v4.3.2). Quality control filtering was applied to remove cells with mitochondrial gene content exceeding 15%. Filtered data were normalized using the SCTransform function within Seurat. Processed datasets from individual samples were merged, and batch effects between experiments were corrected using Harmony (v1.2.1). Integrated data underwent dimensionality reduction (UMAP) and clustering (Leiden algorithm). Clusters were manually annotated based on the expression of established cell-type-specific marker genes (**Table S2**).

### Blood-Brain Barrier Gene Expression Profiling

Gene sets for blood-brain barrier-related biological processes as establishment of BBB [GO:0060856], and maintenance of BBB [GO:0035633]) were retrieved from the Gene Ontology database via biomaRt package (v2.58.2)^79^. Pseudobulk expression values were generated by aggregating SCT-normalized data by cell type and condition, and log2 fold changes between assembloids and control organoids were calculated for each cell type. Heatmaps were generated using ComplexHeatmap package (v2.18.0)^80^. Hierarchical clustering was applied to both genes (rows) and cell types/conditions (columns).

### Blood-Brain Barrier Gene Signature Analysis

To evaluate blood-brain barrier (BBB) characteristics, genes associated with the Gene Ontology (GO) term "Establishment of blood-brain barrier" (GO:0060856) were retrieved using the biomaRt package. UCell (v2.6.2) enrichment scores for this BBB gene signature were computed for each cell in the dataset^81^. Cells were categorized as brain-derived or vascular-derived based on lineage identity. Statistical comparisons of BBB signature scores between organoid groups within each lineage classification were performed using the Wilcoxon rank-sum test. Results were visualized using boxplots with p-values adjusted for multiple comparisons where applicable.

### Endothelial Cell Annotation and Zonation Analysis

Endothelial cells (ECs) were computationally isolated from the organoid dataset for detailed characterization. Sub-clustering and dimensionality reduction were performed on the EC subset to identify distinct subpopulations. To investigate arteriovenous zonation, enrichment scores for arterial, capillary, and venous gene signatures were calculated for each EC subcluster using UCell (v2.6.2). The gene sets used for scoring were curated from published human brain vascular datasets (**Table S3**). EC clusters were annotated based on their dominant zonation signature enrichment profiles. Differences in zonation signature score distributions between organoid types were assessed using the Wilcoxon rank-sum test and visualized as violin plots.

### Gene Ontology Analysis

GO enrichment analysis was conducted to identify biological processes associated with DEGs between fused organoids and other organoid types. For endothelial cells, genes between vascularized cortical assembloids and vascular organoids were identified using the Wilcoxon rank-sum test, applying thresholds of absolute log2 fold change > 0.25 and FDR-adjusted p-value < 0.05. Mitochondrial, and ribosomal genes were excluded. GO enrichment for Biological Process (GO:BP) terms was performed using the gprofiler2 package (v0.2.3), and results were visualized using horizontal bar plots. Selected vascular-related terms were also compared between fused and vascular organoids using grouped bar plots. For the cell type–resolved analysis, differential expression was assessed between vascularized cortical assembloids and a combined reference group of brain and vascular organoids, using a generalized linear model that included organoid condition and sample identity as covariates. Genes with an effect size > 0.15 and FDR-adjusted p-value < 0.005 were considered significant. GO:BP enrichment was conducted separately for upregulated and downregulated genes per cell type using the clusterProfiler package (v4.10.1), with the universe defined as all expressed genes. Enrichment significance for selected developmental and vascular terms was visualized using ComplexHeatmap (v2.18.0), with non-significant values assigned a baseline of zero (-log10 p = 0).

### Differential Expression Analysis with Random Sampling

To assess transcriptional differences between vascularized cortical assembloids (vCAs) and cortical organoids (COs) across specific neural cell populations, we implemented a randomized subsampling strategy followed by differential gene expression analysis. Ten rounds of random sampling with replacement were performed independently for each selected cell type: neuronal progenitor cells (NPC), interneurons (IN), excitatory neurons (EN), mitotic cells, and astrocytes (AC). In each iteration, up to 750 cells were randomly sampled from each condition group (vCA and CO), depending on cell availability.

Differential gene expression analysis was conducted using the MAST implemented in the Seurat (v5.1), with organoid type and iPSC cell lines included as latent variables to account for batch effects. Genes were considered significant if they had an adjusted p-value (Bonferroni-corrected) < 0.01, an absolute average log2 fold change > 1, and were detected in more than 10% of cells in both groups. Genes corresponding to mitochondrial genes (MT-), ribosomal proteins (RPS, RPL), or ENSEMBL gene identifiers (ENSG) were excluded from the results.

The number of significant DEGs identified in each iteration was recorded to quantify transcriptional divergence for each cell type. These values were compared using analysis of variance (ANOVA), followed by Tukey’s Honest Significant Difference (HSD) post-hoc test to determine statistically significant differences in DEG counts across cell populations and conditions.

### CellChat Communication Analysis

Cell to cell communication was inferred using CellChat (v2.1.2) to compare interactions between vascularized cortical assembloids (vCAs) and the combined group of cortical organoids (CO) and vascular organoids (VO). Separate models were constructed for each group based on cell type annotations and RNA assay expression data. Pathway-specific interaction strength was visualized using with chord plots and barplot.

### Cell Cycle Analysis

Tricycle (v1.10.0) was applied to log-normalized expression data to infer continuous cell cycle positions using a human reference. Cells were assigned to discrete phases, and chi-squared tests revealed significant differences in phase distribution across organoid groups for all major cell types. To highlight condition-specific trends relevant to our study, we focused on the M.G1 phase and quantified its proportion per organoid and cell type.

### Assessment of Brain-Enriched Endothelial and Pericyte Cell Signature in Organoids

Organ-specific endothelial and pericyte cell gene expression data were obtained from the supplementary material^51^. The top 25 genes with the highest average log fold change in brain endothelial and pericyte cells compared to other organs were selected to define a brain-specific cell signatures. This gene set was used to compute UCell enrichment scores within respective cell types across all organoid conditions. Scores were calculated using the default UCell workflow in R and visualized as violin plots to compare enrichment across groups.

### Primary Tissue Sequencing Data Integration and Processing

To benchmark organoid cell populations against their *in vivo* counterparts, published scRNA-seq datasets from developing human brain vasculature^11,12^ were integrated with our organoid dataset. Primary tissue datasets were preprocessed according to their original publication methods, including quality control and normalization. Batch correction and data integration were performed using scVI (v1.1.2), followed by semi-supervised cell type annotation refinement using SCANVI (v1.1.2) trained on established marker genes to ensure accurate alignment of shared populations.

### Organoid Protocol Comparison

To assess the transcriptional similarity between brain organoids and primary human brain tissue, we calculated cosine similarity scores across major cell types using pseudobulk gene expression profiles. For each dataset, pseudobulk profiles were generated by averaging log-transformed expression values across all cells within a given cell type, using count-based normalization to ensure comparability. Cell type–specific gene signatures were defined by selecting the top 1,000 differentially expressed genes from the primary tissue dataset based on statistical significance and effect size thresholds. Cosine similarity was then computed between primary tissue profiles and those derived from each organoid protocol^25,27-29^. This approach is consistent with previously published methods for benchmarking transcriptional fidelity across platforms and experimental conditions^70-72^.

### BBB Gene Signature Scoring

Endothelial cells from vascular cerebral assembloids (vCA) generated using our protocol were compared to endothelial cells from other published protocols^25,27-29^. Cells were scored for expression of blood-brain barrier-related gene signatures with related gene ontology terms using UCell(v2.6.2)^81^. Statistical significance was assessed using Wilcoxon rank-sum test with Bonferroni correction.

## Extended Data Figures Figure Legends

**Extended Data Fig. 1 Reproducibility and fidelity of vascular network formation across hiPSC lines a**, Representative whole mount confocal microscopy analysis showing CD31^+^ endothelial cells (cyan) in day 8 sprouting vascular organoids. DAPI (white) labels cell nuclei. Lower magnification scale bar, 150 µm. Inset, high magnification view. Arrowheads, CD31^+^ vascular sprouts. Scale bar, 100 µm. **b**, Representative confocal microscopy analysis of CD31^+^ endothelial networks in primary human cortical plate tissue and vascularized cortical assembloids (vCAs). GW, gestational week. Scale bars, 100 µm. **c**, Quantitative vascular morphometry of branching index (left), vascular density (middle), and total vessel length (right) in three hiPSC cell lines (red) and human primary cortical plate tissue. n = 3 organoids per cell line; n = 4 donors for primary tissue samples. ns, non-significant.

**Extended Data Fig. 2 Quality control metrics for scRNA-sequencing. a.** Violin plots showing number of genes (left), unique molecular identifiers (middle), and mitochondrial percentage (right) across cell populations. EC, endothelial cell; PC, pericyte; FB, perivascular fibroblast; MG, microglia; Mitotic, mitotic cell; NPC, neural progenitor cells; RG, radial glia; AC, astrocyte; IN, interneuron; EN, excitatory neuron. **b-c**, Bar graph of cell proportion by organoid type (B) or hiPSC cell line (C). vCA, vascularized cortical assembloids; VO, vascular organoid; CO, cortical organoid.

**Extended Data Fig. 3 Blood-brain barrier signatures and neurovascular cell-to-cell communication pathways in vascularized cortical assembloids. a,** Boxplots of establishment of blood-brain barrier gene enrichment scores across. VO, vascular organoid; vCA, vascularized cortical assembloids; CO, cortical organoids. ***, *P* < 0.001. **b-c**, Violin plots showing brain organotypic endothelial (B) and pericyte (C) cell signatures in VOs (gray) and vCAs (red). ***, *P* < 0.001. **d**, Cell-to-cell communication pathways across cell populations ranked on their differences of overall information flow within inferred networks in vCAs (red) or *in silico* assembled controls (orange, VO + CO). Red text, pathways significantly overrepresented in vCAs; orange text, pathways statistically overrepresented in controls; black text, not significant. **e,** Chord plot of statistically enriched incoming endothelial pathways in vCAs. **f**, Chord plot showing incoming Wnt and non-canonical Wnt signaling to endothelial cells in vCAs. Arrow thickness is proportional to interaction strength. **g**, Chord plot showing outgoing endothelial pericyte recruitment pathways in vCAs. Arrow thickness is proportional to interaction strength. **h,** Bar plots showing proportion of M to G1 cell cycle proportions for endothelial(blue), fibroblast(beige), pericytes(green). ***, *P* < 0.001.

**Extended Data Fig. 4 Vascularization promotes neurodevelopment in vascularized cortical assembloids. a,** Heatmap visualization of enriched neurodevelopmental gene ontology pathways in vascular cells within vascularized cortical assembloids (vCAs). Gray, low enrichment; Red, high enrichment. **b**, Volcano plot visualization of differentially expressed genes in neural progenitor cells in vCAs. Wilcoxon rank sum, FDR corrected p-value < 0.01 and log_2_FC > 1.0 to be colored significant. Red, upregulated; Blue, downregulated. **c**, Representative confocal microscopy analysis showing DCX^+^ neuroblasts in cortical organoids (COs) and vCAs. DAPI (white) labels cell nuclei. Scale bar, 100 µm. **d**, Representative confocal microscopy analysis showing SYP^+^ pre-synaptic (red) and HOMER^+^ post-synaptic (cyan) densities in COs and vCAs. DAPI (white) labels cell nuclei. Right, quantification of SYP^+^ and Homer^+^ synapses. n = 3 hiPSC lines per condition, 3-5 organoids per cell line per condition, all replicates shown. **, *P* < 0.01. Scale bar, 10 µm.

**Extended Data Fig. 5 Atlas-level integration quality control metrics. a,** Violin plots showing number of genes (left), unique molecular identifiers (middle), and mitochondrial percentage (right) across cell populations after scVI integration. EC, endothelial cell; PC, pericyte; FB, perivascular fibroblast; MG, microglia; Mitotic, mitotic cell; NPC, neural progenitor cells; RG, radial glia; AC, astrocyte; IN, interneuron; EN, excitatory neuron. **b**, Sankey plots showing harmonization of cell annotations across published vascularization protocols with available scRNA-seq data. Line width is proportional to cell numbers.

## Code & Data Availability

The scRNA-seq dataset supporting results of this paper will be deposited at GEO and SRA upon publication. This paper does not report original code.

## Schematics & Figures

Schematics and figures were created with BioRender and Adobe Illustrator 2025.

## Acknowledgements

We are grateful to Vinh Nguyen, and the Flow Cytometry Colab at the University of California, San Francisco for their consultation and support during the Fluorescence-activated cell sorting (FACS). We also acknowledge the Hervey-Jumper Lab for providing equipment access for confocal microscopy, and calcium imaging assays. We further thank Gemma K. Alderton for her manuscript revisions. This work has been supported by the Shuri and Kay Curci Foundation Award, Marcus Precision Medicine Grant, and Cerebrovascular Section / Congress of Neurological Surgeons Foundation Young Investigators Research Grant (to E.A.W).

## Author Contributions

J.M.R and E.A.W conceptualized the study. M.Ö., and J.M.R. provided and organized iPSC cell lines and organoids, J.M.R. and M.Ö. performed FACS-enrichment and scRNA sequencing of organoids. B.G., J.C.W, and C.N.K performed computational analyses. D.S, S.B., M.Ö., S.A., A.G., M.Ö., and J.M.R. performed immunostaining, imaging and quantifications of organoids. S.O., and D.S. performed calcium imaging and quantifications. B.G., D.S., S.B., T.M. prepared schematics and figures. T.M., D.S., B.G., S.B., and J.C.W. prepared supplementary tables. E.A.W. provided laboratory space, computational workstations, and funding acquisitions. B.G., S.B., and E.A.W. wrote the manuscript with input from all authors. D.S., J.M.R., S.A., A.G., M.Ö., T.M., J.A., J.C.W, J.K., C. N. K., Y.R., S.W., H.K., E.E.C., E.F.C, D.H., M.T., S.O., A.R.K and T.J.N. assisted with data interpretation. E.A.W. supervised the study.

## Materials & Correspondence

Requests for materials or correspondence should be directed to Ethan A. Winkler, MD, PhD.

## Supplementary Tables

Supplementary Table 1: Clinical metadata of iPSC cell lines

Supplementary Table 2: Cell type marker genes

Supplementary Table 3: Endothelial cell arteriovenous specification markers

Supplementary Table 4: Comparison of published organoid vascularization protocols

