## Supplementary figures and images for "Scalable high-fidelity human vascularized cortical assembloids recapitulate neurovascular co-development and cell specialization"

### Extended Data Figures

##### Extended Data Fig. 1

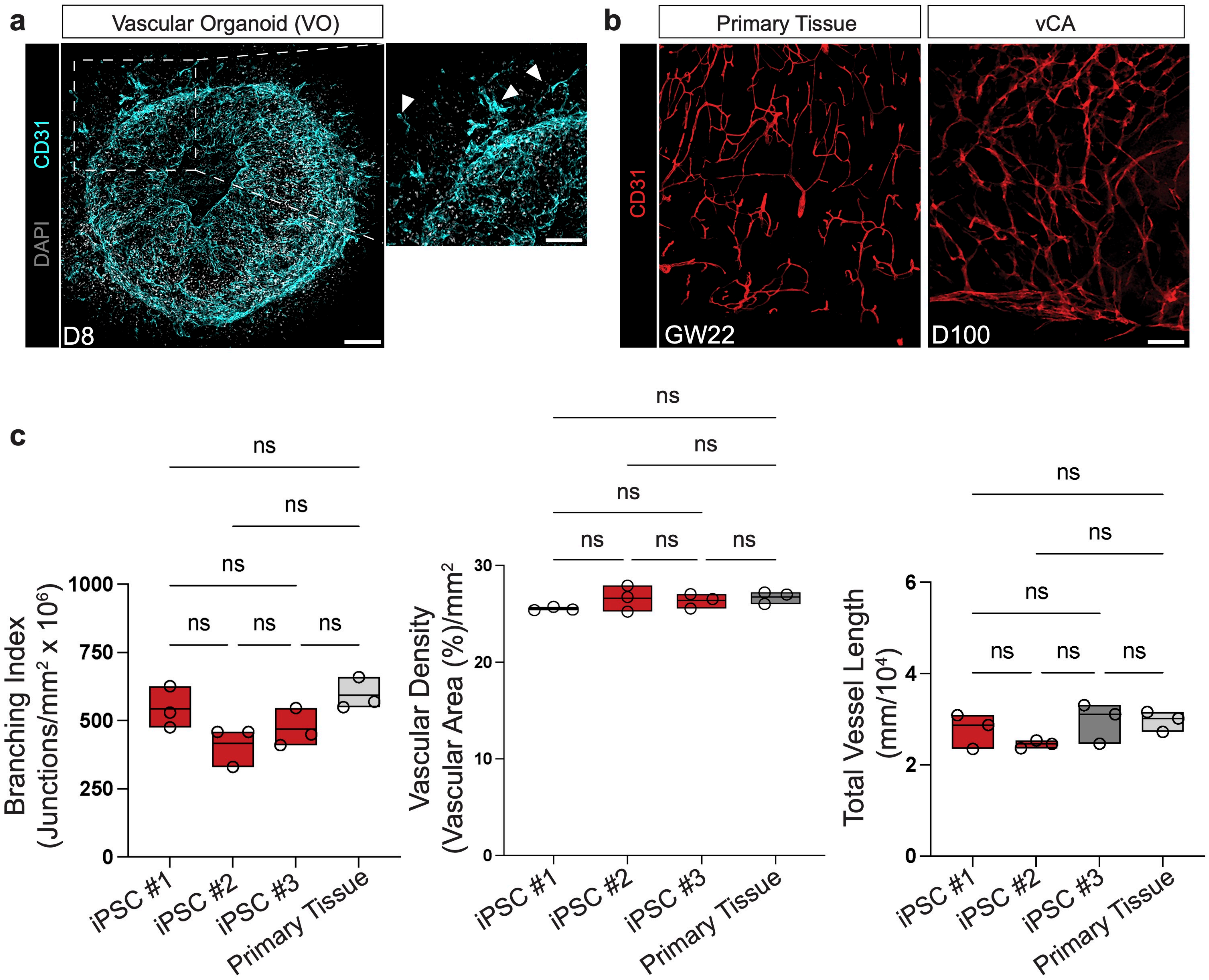

#### Extended Data Fig. 2

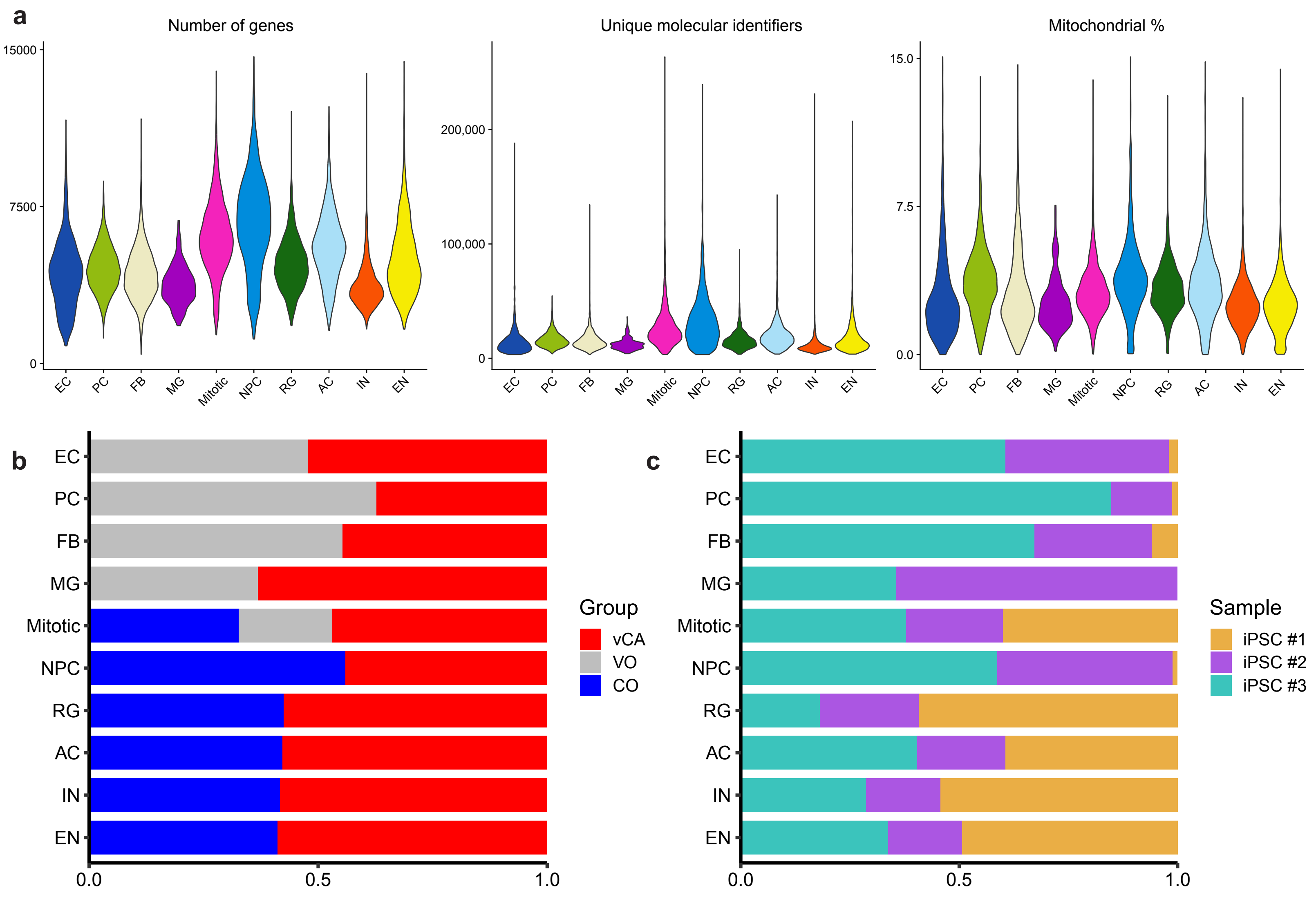

#### Extended Data Fig. 3

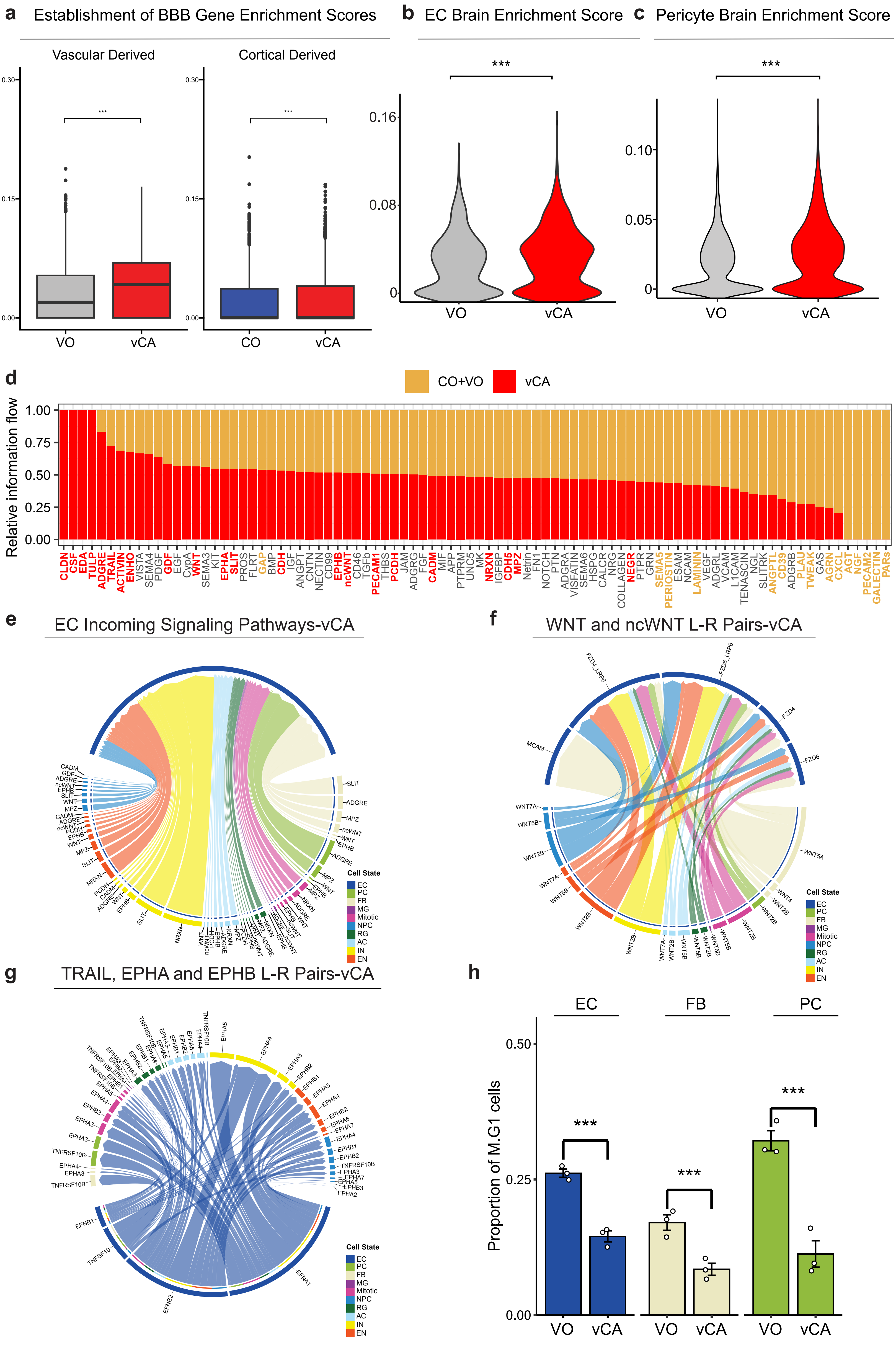

#### Extended Data Fig. 4

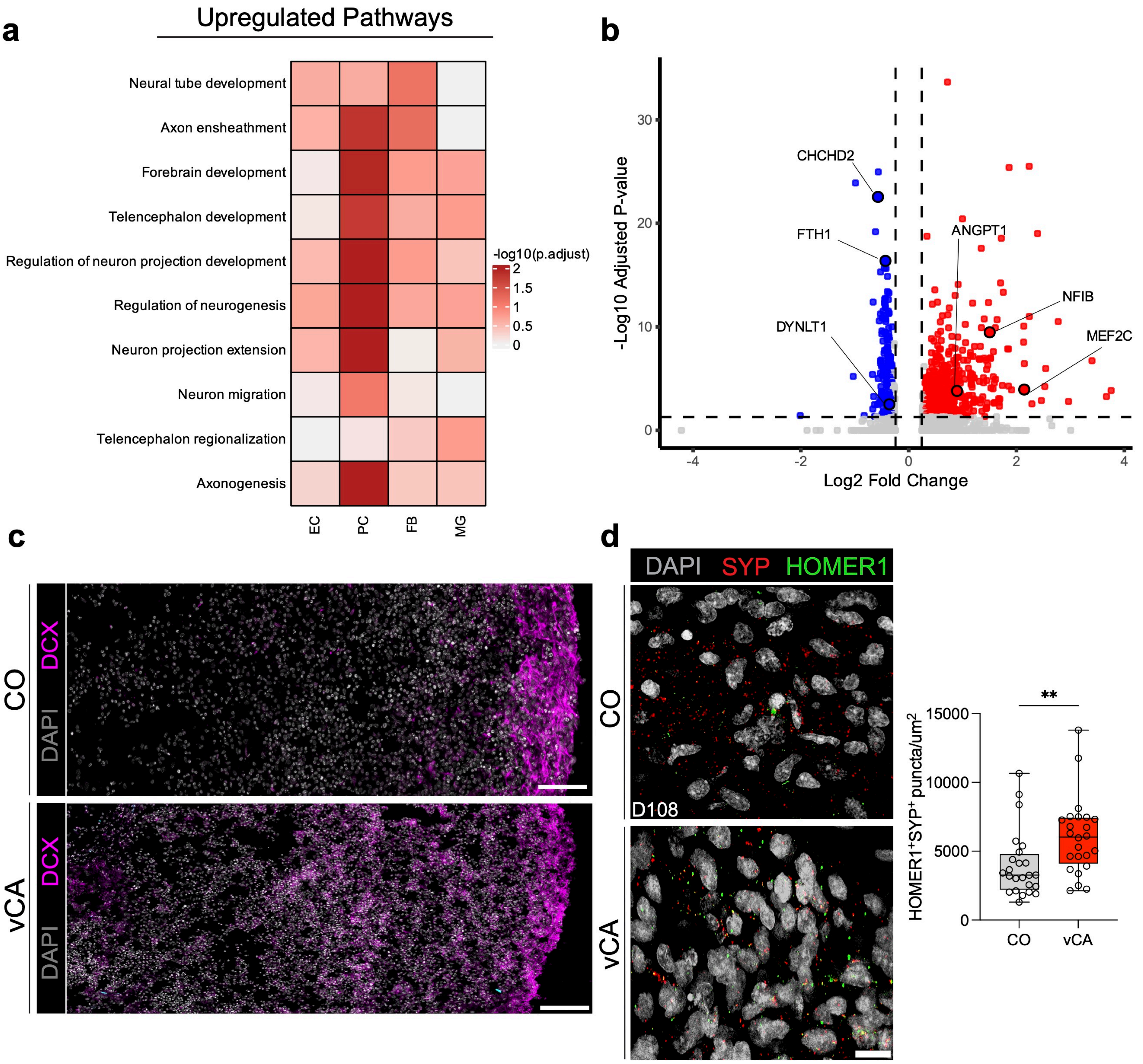

#### Extended Data Fig. 5

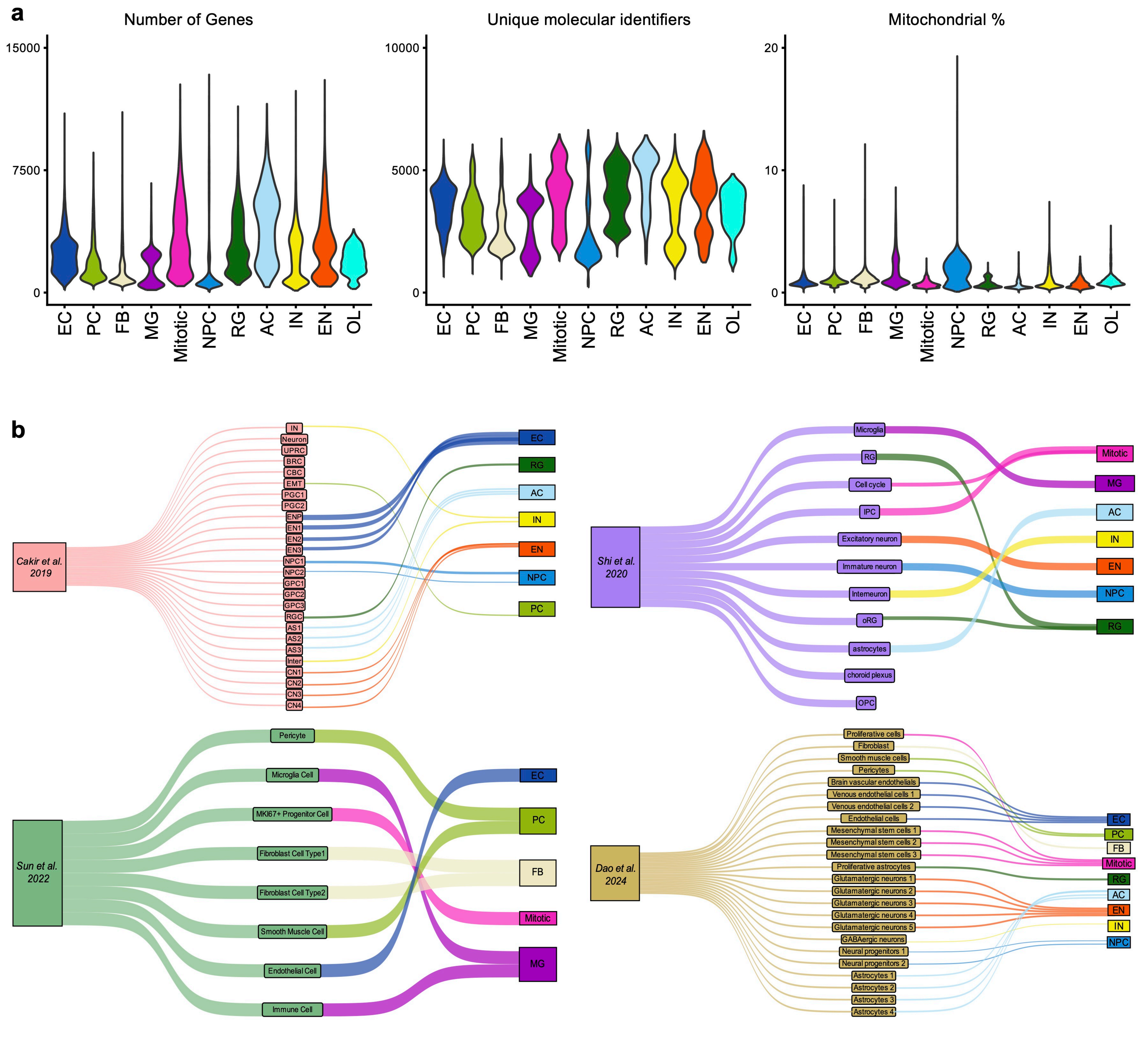
